# RGG-motif protein Sbp1 is required for Processing body (P-body) disassembly

**DOI:** 10.1101/2021.02.23.432385

**Authors:** Raju Roy, Ishwarya Achappa Kuttanda, Nupur Bhatter, Purusharth I Rajyaguru

## Abstract

RNA granules are conserved mRNP complexes that play an important role in determining mRNA fate by affecting translation repression and mRNA decay. Processing bodies (P-bodies) harbor enzymes responsible for mRNA decay and proteins involved in modulating translation. Although many proteins have been identified to play a role in P-body assembly, a bonafide disassembly factor remains unknown. In this report, we identify RGG-motif translation repressor protein Sbp1 as a disassembly factor of P-bodies. Disassembly of Edc3 granules but not the Pab1 granules (a conserved stress granule marker) that arise upon sodium azide and glucose deprivation stress are defective in *Δsbp1*. Disassembly of other P-body proteins such as Dhh1 and Scd6 is also defective in *Δsbp1*. Complementation experiments suggest that the wild type Sbp1 but not an RGG-motif deletion mutant rescues the Edc3 granule disassembly defect in *Δsbp1*. We observe that purified Edc3 forms assemblies, which is promoted by the presence of RNA and NADH. Strikingly, addition of purified Sbp1 leads to significantly decreased Edc3 assemblies. Although low complexity sequences have been in general implicated in assembly, our results reveal the role of RGG-motif (a low-complexity sequence) in the disassembly of P-bodies.

## Introduction

RNA granules are conserved, membraneless dynamic condensates consisting of RNA-protein complexes. They play an important role in regulation of mRNA translation and decay to modulate cell proteome in response to specific physiological cues. P-bodies (PB) and stress granules (SG) are the most well-characterized form of RNA granules. Stress granule and P-bodies share many protein components and exchange mRNPs (Brengues and Parker, 2007; Hoyle et al., 2007; Kedersha et al., 2005). SGs harbor mRNAs in translationally repressed forms in complex with various translation factors (Buchan et al., 2008; Kedersha et al., 1999; Mazroui, 2006). On the other hand, PBs also contain enzymes and proteins that promote mRNA decay thereby leading to the degradation of resident mRNAs (Fenger-Gron et al., 2005; Parker and Sheth, 2007). Interestingly in yeast, SG arise from pre-formed PB indicating that PB acts as a repository of mRNPs that are either degraded or transferred to SG subsequently (Buchan et al., 2008).

PBs exist at a basal level under unstressed conditions in some cell types including yeast and are strongly induced in response to several stresses such as glucose starvation, oxidative stress, heat stress which lead to global translation repression (Buchan et al., 2008). A recent report indicates that PBs are important for maintaining stem cell plasticity as the loss of PBs leads to a differentiation-resistant state of the embryonic stem cells (Di Stefano et al., 2019). Some PB resident proteins regulate mRNAs involved in inflammatory response (Ohn and Anderson, 2010). Virus infections specifically by RNA viruses lead to disassembly of PB indicating that PB could play a role in viral stress response. PBs have been reported to contain mRNA encoding proteins involved in chromatin regulation, protein degradation, RNA processing and protein synthesis (Standart and Weil, 2018). Analysis of PB-resident mRNAs provides a striking observation. Transcripts encoding regulators of important processes such as protein turnover, RNA processing, cell cycle and energy metabolism are compartmentalized to PB however the transcripts that encode proteins that constitute the respective machineries are excluded (Standart and Weil, 2018). Such arrangement highlights the role of PB in specifically regulating the regulators. Therefore, understanding PB assembly and disassembly would be crucial to elucidating regulation of important cellular processes.

The mechanistic basis of PB assembly has been addressed. An important factor that modulates RNA granule formation in general is intrinsically disordered regions (IDRs) in RNA granules associated proteins (Protter and Parker, 2016). IDRs often contain repeats of specific amino acid residues (low complexity sequences) that cannot form a three-dimensional structure but can be involved in binding proteins and/or nucleic acids to contribute specific functions (van der Lee et al., 2014). Some examples of low-complexity (LC) sequences include repeats of RGG/RG (Arg-Gly-Gly), YGG (Tyr-Gly-Gly), QQQ (or, polyQ), GY/GSYGS/ GYS/SYG/ SYS, Q/N (Gln-Asn), RS (Arg-Ser) and PPP (Proline-rich motif). These LC regions containing protein are known to bind to other proteins, and in specific cases, it can bind to self, leading to the formation of higher-order structures (Calabretta and Richard, 2015). Edc3 is a conserved marker of PB that plays a vital role in its assembly. The C-terminal Yjef-N domain if Edc3 potein binds itself to promote formation of higher order PB assembly (Decker et al., 2007). Deletion of Edc3 reduces the ability of cells to form PB however *Δedc3* cells are highly defective in PB assembly in the absence of the C-terminal Q/N-rich prion-like motif of PB protein Lsm4 (Decker et al., 2007). Consistent with the idea of Edc3 self-association in yeast other PB proteins are known to multimerize and contribute to PB assembly. Human EDC3 forms dimers (Ling et al., 2008), fly DCP1 forms trimers (Tritschler et al., 2009) and human RCK/p54 can form multimers (Ernoult-Lange et al., 2012). Similarly PGL-3, a key germ granule assembly factor in *C. elegans* self-associates and binds other RNA and mRNP components through its RGG-motif (Hanazawa et al., 2011). TIA-1, an essential stress granule protein in humans facilitates SG assembly via aggregation of the Q/N rich region (Gilks et al., 2004; Kawakami et al., 1992).

The LC sequences participate in multiple low affinity interactions to undergo liquid-liquid phase separation (LLPS) that promotes RNA granule formation (Banani et al., 2016; Kato et al., 2012). RNA often provides a scaffold for such interactions between the RNA binding proteins with LC sequences to augment LLPS (Fernandes and Buchan, 2020). A strong link has emerged between aberrant phase transitions of RNA granule components and neurodegenerative disorders. FUS and TDP43 (RGG-motif containing nuclear proteins involved in amyloid lateral sclerosis) are reported to accumulate mutations that lead to formation of cytoplasmic amyloid inclusions which are linked to the disease (Patel et al., 2015; Pesiridis et al., 2009). Similar observations are also reported for other nuclear RBPs such as hnRNPA1, hnRNPA2 and EWSR1 (Couthouis et al., 2012; Kim et al., 2013). These examples highlight that persistant aberrant RNA granules that fail to disassemble play an important role in these neurodegenerative disorders (Advani and Ivanov, 2020). Therefore understanding factors that contribute to disassembly is paramount as this information could help device strategies to deal with disassembly-resistant RNA granules. In this study, we report that RGG-motif protein Sbp1 is required for P-body disassembly and RGG-motif of Sbp1 plays an important role in promoting disassembly.

## Results

### *Δsbp1* is defective in Edc3 granule disassembly

To test the role of RGG-motif proteins in RNA granule disassembly, we took strains of *Saccharomyces cerevisiae* (BY4741) where the known translation repressors, Scd6 (Poornima et al., 2016; Rajyaguru et al., 2012) and Sbp1 (Bhatter et al., 2019; Brandariz-Núñez et al., 2018) is deleted, respectively. A *CEN* plasmid co-expressing Pab1-GFP (a stress granule marker) (Buchan et al., 2011) and Edc3-mCherry (a P-body marker) (Buchan et al., 2011) was transformed into wild-type, *Δscd6* and *Δsbp1* strains, respectively. Wild-type strain upon sodium azide stress showed induction of stress granule (SG) and P-body (PB) formation consistent with earlier reports (Buchan et al., 2011). These granules disassembled upon removal of stress during recovery (Figure 1A, wild-type microscopy panel; Figure 1B). *Δscd6* strain showed reduced stress granule formation, as reported earlier (Rajyaguru et al., 2012), but P-body assembly was comparable to the wild-type strain (Figure 1A, *Δscd6* microscopy panel; Figure 1B). *Δsbp1* strain also showed defect in stress granule formation but normal P-body formation as compared to wild-type strain (Figure 1A, *Δsbp1* microscopy panel, stressed condition, Figure 1B). A striking observation was that *Δsbp1* was defective in P-body disassembly after 1h recovery process (Figure 1A, *Δsbp1* microscopy panel, recovery condition; Figure 1B). The disassembly defect of Edc3-mCherry in *Δsbp1* strain is not due to increased levels of Edc3-mCherry protein during recovery (Figure 1C). The disassembly of stress granules was found to be normal (Figure 1A, *Δsbp1* microscopy panel, recovery condition). Based on these results, we conclude that disassembly of Edc3 foci is defective in *Δsbp1* strain.

**Figure 1:**
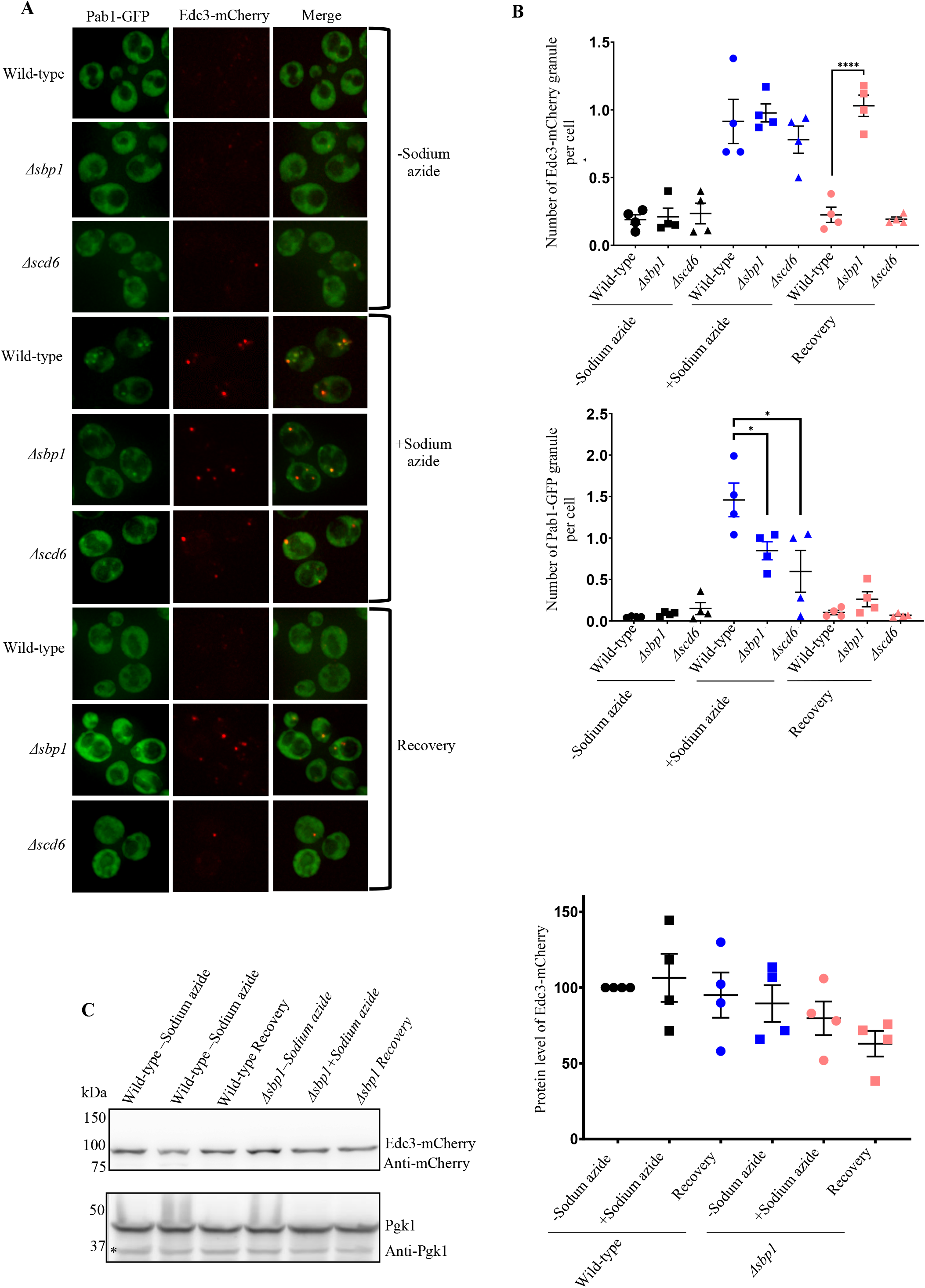
Disassembly of Edc3 granules is defective in *Δsbp1*. A) *Δsbp1*leads to Edc3 granule disassembly defect. Cells were cultured till 0.35-0.4 O.D. and incubated for 30 minutes with or without 0.5% (v/v) Sodium azide at 30°C. Susequently, cells were pelleted by centrifugation (4200g, 10 seconds, RT) and washed thrice with glucose containing medium. For stress recovery, the resuspended cells were grown for an additional 1 hour at 30°C in media without Sodium azide. B) Graph depicting the number foci per cell in wild-type, *Δsbp1*and *ΔSCD6* strain in various culture conditions (p<0.005). C) Protein levels of Edc3-mCherry during conditions shown in A.

### Disassembly of other P-body markers is defective in *Δsbp1*

We next tested if *Δsbp1* affects disassembly of other P-body components such as Dhh1 and Scd6. Dhh1 and Scd6 are PB components (Nissan et al., 2010) that are important for P-body assembly. We transformed wild-type and *Δsbp1* strains with *CEN* plasmid expressing either Dhh1-GFP or Scd6-mCherry. The assembly of Dhh1-GFP granules was comparable to that of wild type cells however the disassembly of Dhh1-GFP granules was significantly defective during recovery in *Δsbp1* strain as compared to wild-type (Figure 2A & B). The disassembly defect was not due to increased protein levels of Dhh1-GFP (Figure 2C). Similarly, the disassembly but not the assembly of Scd6-mCherry granules was affected in *Δsbp1* (Figure 2D & E). The protein level of Scd6-mCherry remained unaffected in recovery as compared to stress conditions (Figure 2F). Based on these observations we conclude that PB disassembly is defective in *Δsbp1*.

**Figure 2.**
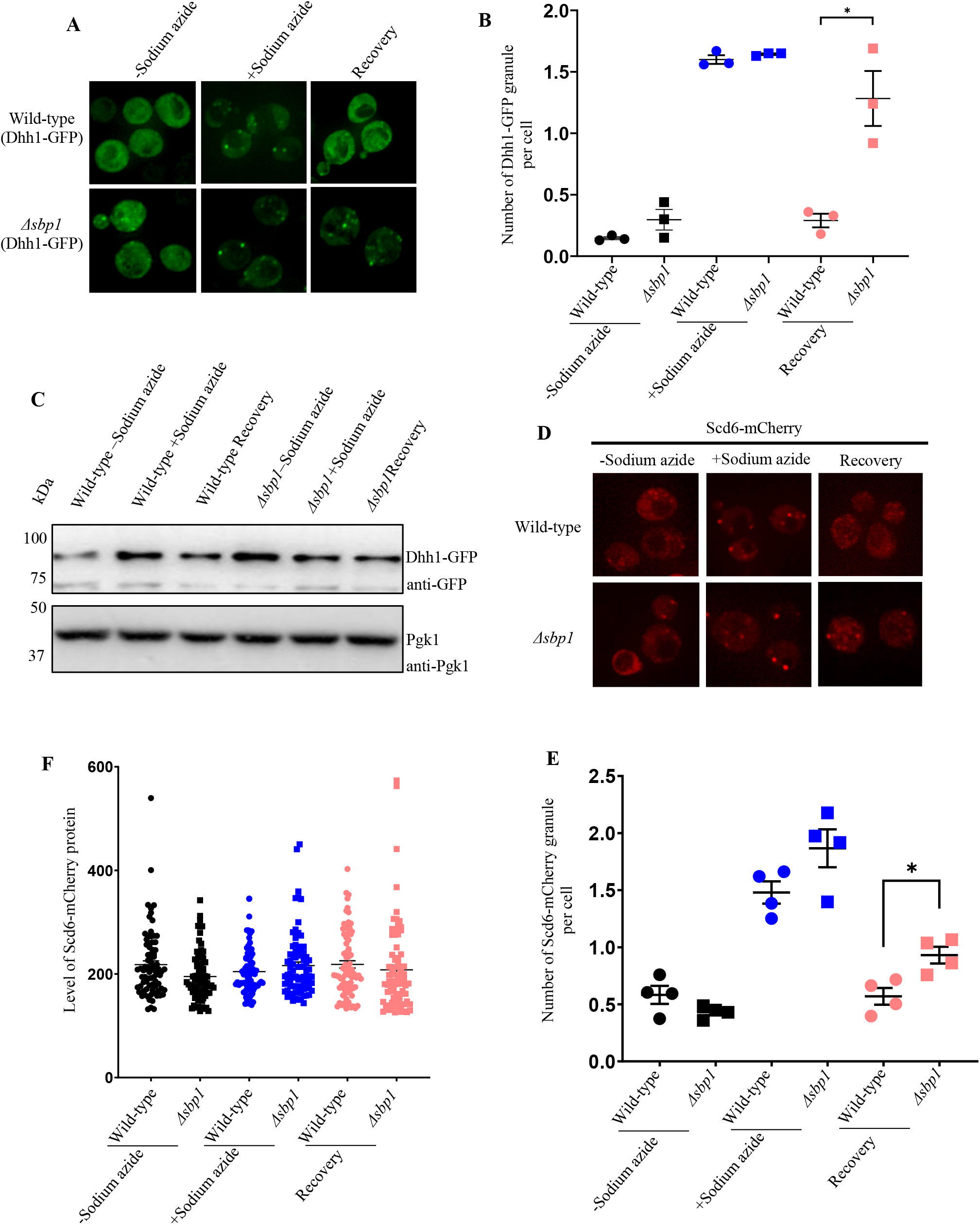
Disassembly of other P-body markers Scd6 and Dhh1 is defective in absence of Sbp1. A) Live cell microscopy for the recovery experiment using Scd6-mCherry as P-body marker. B) Graph depicting the number of Scd6-mCherry foci per cell in wild-type and *Δsbp1* strain (p<0.05). C) Protein level of Dhh1-GFP during unstress, stress and recovery in wild-type and *Δsbp1* strain. D) Live cell microscopy for the recovery experiment using Dhh1-GFP as P-body marker. E) Graph depicting the number of Dhh1-GFP foci per cell in wild-type and *Δsbp1*strain (p<0.05). F) Protein level of Dhh1-GFP during unstress, stress and recovery in BY4741 and *Δsbp1* strain.

### Edc3 granule disassembly defect in *Δsbp1* is not stress-specific

The dynamics and composition of RNA granules differ between stress conditions (Buchan et al., 2011). Therefore, we tested whether P-body disassembly defect in *Δsbp1* strain was stress-specific. We analyzed disassembly of granules that are induced upon glucose deprivation in *Δsbp1*. Glucose deprivation leads to global translation repression and consequently RNA granule assembly (Brengues et al., 2005). As expected, the WT strain showed normal assembly and disassembly of stress granules (Pab1-GFP) and P-bodies (Edc3-mCherry) as reported earlier (Buchan et al., 2008) (Figure 3A; wild-type microscopy panel; Figure 3B). Assembly of stress granules and P-bodies was comparable to that of wild-type in *Δsbp1* strain (Figure 3A; *Δsbp1* minus glucose panel; Figure 3B). Interestingly, the disassembly of P-bodies and not the stress granules was defective in *Δsbp1* (Figure 3A; *Δsbp1* recovery panel; Figure 3B). Protein levels of Edc3-mCherry during recovery were comparable to those under stress conditions (Supplementary Figure 1). This suggests that the Sbp1 affects P-body disassembly under multiple stress conditions.

**Figure 3.**
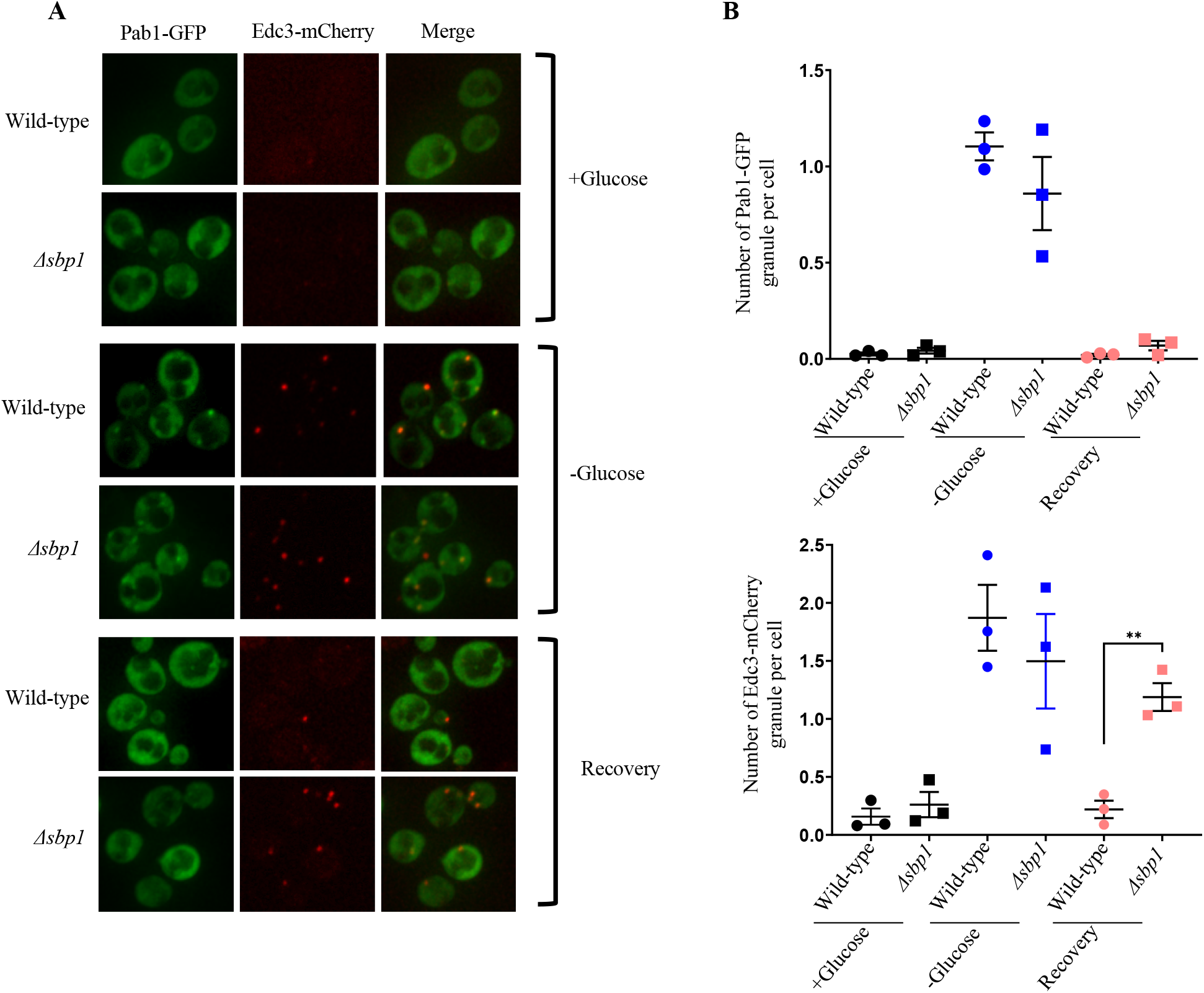
Edc3-mCherry granule disassembly defect is not stress specific. Live-cell microscopy for the recovery experiment following glucose deprivation as stress condition. (A)Live cell imaging of wild-type strain and *Δsbp1*strain upon glucose starvation. (B) Graph depicting the number of foci per cell in wild-type and *Δsbp1*strain in various culture conditions during granule recovery experiment.

### RGG-motif of Sbp1 is required for rescuing Edc3 granule disassembly defect in *Δsbp1*

To confirm if the disassembly defect observed in *Δsbp1* was indeed mediated by Sbp1, we complemented *Δsbp1* strain with CEN plasmid (pRS315) expressing wild-type Sbp1 (pRS315-*SBP1*). The *Δsbp1* strain transformed with empty-vector was defective in both Pab1-GFP assembly (Supplementary Figure 2A; *Δsbp1*-pRS315(EV); 2B) and Edc3-mCherry disassembly (Figure 4B & C, *Δsbp1*-pRS315(EV) recovery panel) as compared to the wild-type empty-vector strain (Figure 4B & C, wild-type-pRS315(EV) recovery panel). We observe that the *Δsbp1* strain transformed with pRS315(*SBP1*) showed disassembly of Edc3 granule formed during stress comparable to the wild type cells (Figure 4B & C; *Δsbp1*-pRS315(*SBP1*) recovery panel). We conclude that disassembly defect in the *Δsbp1* strain is indeed due to absence of Sbp1 and can be rescued by expressing Sbp1 on a plasmid.

**Figure 4.**
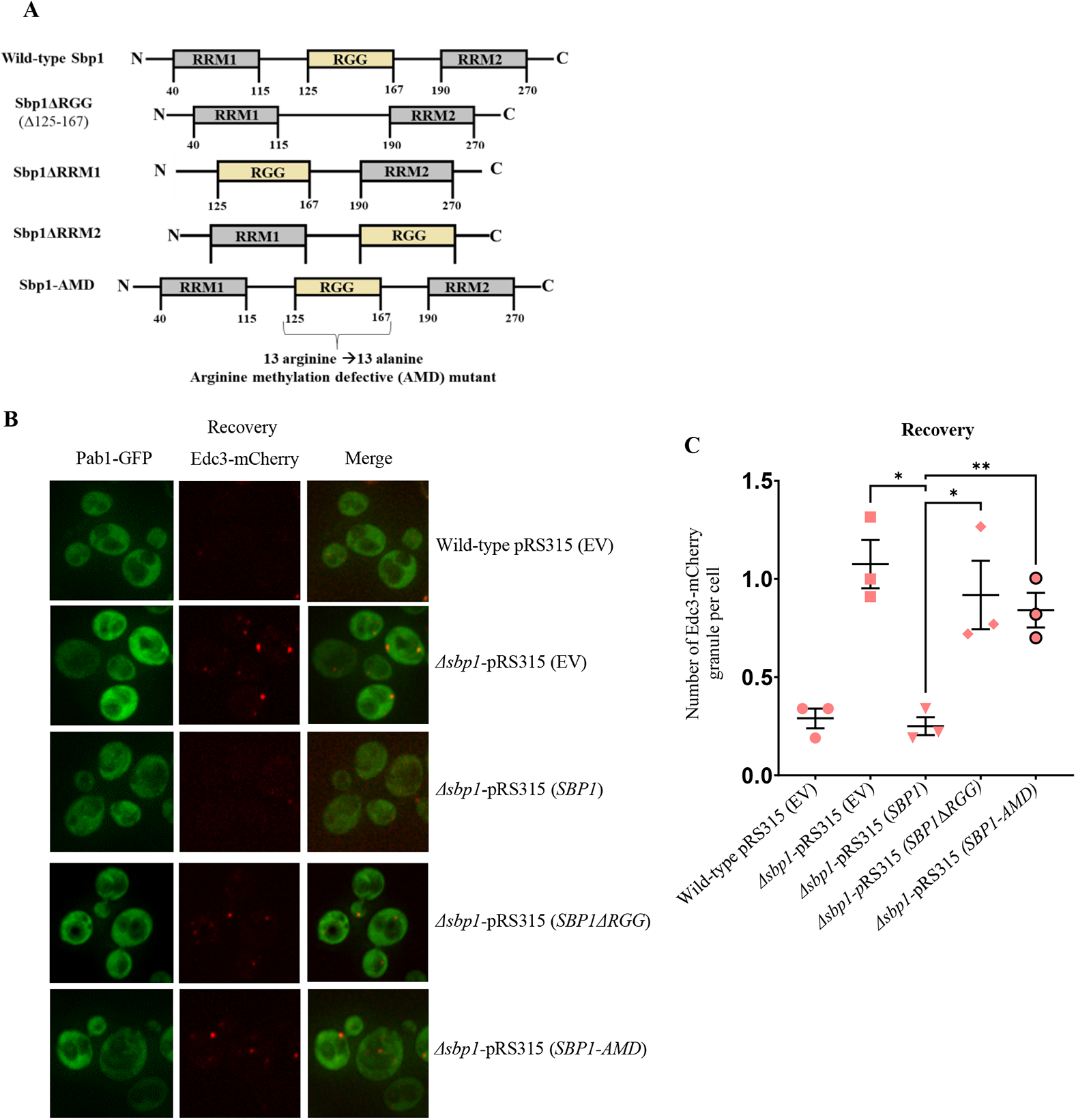
Complementation of *Δsbp1* with wild-type Sbp1 and Sbp1 mutants. A) Domain organization of Sbp1 and mutants used for the study. B) Live-cell microscopy for the recovery experiment following Sodium azide stress condition. Plasmid containing wild-type Sbp1 and Sbp1 mutant were transformed in wild-type and *Δsbp1* cells as indicated. Cells were cultured till 0.35-0.4 O.D. and incubated for 30 minutes with or without 0.5% (v/v) Sodium azide at 30°C. Subsequently, cells were pelleted by centrifugation (4200g, 10 seconds, RT) and washed thrice with glucose containing medium. For stress recovery, the resuspended cells were grown for an additional 1 hour at 30°C in media without Sodium azide. C) Quantitation of the complementation experiment in described in B. D) Protein levels of Edc3-mCherry, Sbp1, Sbp1 mutants and PGK1 during recovery.

Sbp1 is a modular protein with RGG-motif present in between two RRM domains (Figure 4A). We next tested the domain of Sbp1 that might be necessary for complementing the Edc3 granule disassembly defect. We created mutants of Sbp1 that were individually deleted for RGG, RRM1 and RRM2 domain in the pRS315(*SBP1*) plasmid (Figure 4A). The *SBP1ΔRGG* mutant was defective in complementing Edc3 granule disassembly in (Figure 4B & C; *Δsbp1*-pRS315(*SBP1ΔRGG*) recovery panel) as compared to to the full-length Sbp1 expressing in *Δsbp1* strain. This mutant was also defective in complementing stress granule assembly defect (Supplementary Figure 2A and B). The *SBP1ΔRRM1* mutant but not the *SBP1ΔRRM2* mutant was defective in complementing stress granule assembly defect (Supplementary Figure 2C and 2D) however both these mutants did not show any defect in assembly and disassembly of Edc3 granules (Supplementary Figure 2C and 2D). This result suggests that the RGG-motif of Sbp1 protein is required for the disassembly process of Edc3 granules.

### Arginine methylation defective (AMD) mutant of Sbp1 fails to rescue the Edc3 RNA granule disassembly defect

Recent studies have demonstrated the importance of arginine methylation in RNA granule assembly and disassembly (Huang et al., 2018; Tsai et al., 2016; Yamaguchi and Kitajo, 2012). Our previous reports suggest that arginine methylation within RGG-motif promotes Sbp1 repression activity (Bhatter et al., 2019). To test whether arginine methylation affects Sbp1 role in Edc3 granule disassembly, we substituted all 13 arginines in the RGG-motif of Sbp1 to alanines (Figure 4A). This construct was defective in methylation and hence was referred to as arginine-methylation defective (AMD) mutant (Bhatter et al., 2019). We observed that the AMD mutant failed to rescue disassembly defect of Edc3 granule after recovery in *Δsbp1* strain (Figure 4B & C; *Δsbp1*-pRS315(*SBP1-AMD*) recovery panel) as compared to the *Δsbp1* strain complemented with wild type *SBP1* gene. We conclude based on this result that the AMD mutant is compromised in complementing Edc3 granule disassembly defect. Hmt1, a major methyltransferases in yeast is known to methylate Sbp1 (Bhatter et al., 2019). To further test the role of arginine methylation in Edc3 granule disassembly, we assessed disassembly of Edc3 granules in *Δhmt1* strain. Interestingly, the disassembly of Edc3 granules in *Δhmt1* was comparable to the same in the wild type strain (Supplementary Figure 3A & 3B).

### Purified Sbp1 binds to LSm and Yjef-N domains of Edc3

Our results indicate a role of Sbp1 in Edc3 granule disassembly, which is dependent on RGG-motif. We hypothesized that physical interaction between Sbp1 and Edc3 could enable the exit of Edc3 from granules during recovery. Owing to the technical difficulties associated with purifying recombinant full-length Edc3 in soluble form, we tested the interaction of recombinant His-Sbp1-FLAG with Edc3 domains such as LSm-FDF-GST, FDF-GST, and Yjef-N-GST. These domains have previously been used for assessing interaction of Edc3 with different PB components (Nissan et al., 2010). Interaction studies revealed that purified Sbp1 bound to both LSm-FDF and Yjef-N domains but not the FDF domain alone. This interaction was independent of RNA as RNAse was added during protein purification as well as during the protein-protein interaction assay (Figure 5). This result suggests that Sbp1 directly interacts with the Lsm and Yjef-N domains of Edc3.

**Figure 5.**
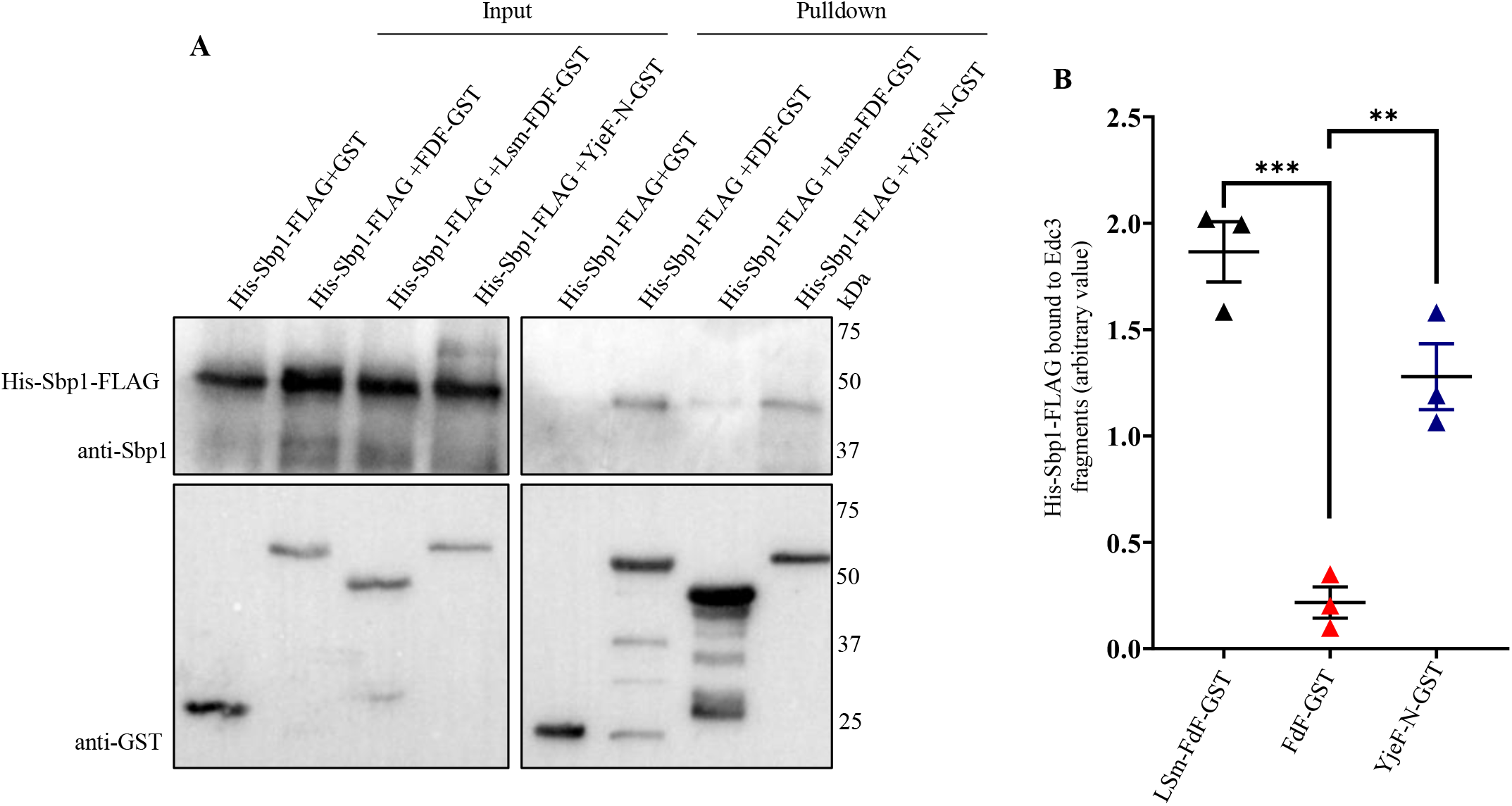
Sbp1 interacts with LSm and YjeF-N domain of Edc3. A. Western blot depicting interaction of Sbp1 with various fragments of Edc3 in presence of RNase A *in vitro*. His-Sbp1-FLAG was purified using Ni-NTA chromatography. Edc3 fragments tagged with GST were purified and immobilized to the glutathione beads. Interaction studies was carried out in 1X protein-protein interaction buffer (PPIB), washed with 1X PPIB post binding and then western was carried out. B. Quantitation depicting level of Sbp1 binds to the fragments of Edc3 in presence of RNase A *in vitro*.

### Sbp1 is required for the disassembly of Edc3-mCherry assemblies *in vitro*

Liquid-liquid phase separation (LLPS) plays an important role in assembly of RNA granules (Wheeler et al., 2016). Proteins with low complexity sequences including Edc3 have been shown to undergo LLPS (Babinchak et al., 2019; Lin et al., 2015) which contributes to the assembly of RNA granules (Damman et al., 2019). Although Edc3 from *S. pombe* has been recently studied to undergo LLPS, Edc3 from *S. cerevisiae* has not yet been reported to form liquid droplets. Therefore, we tested if purified Edc3 underwent phase separation to form higher order assembly. We observed that the purified Edc3-mCherry phase-separated to form small assemblies (Figure 6A). Strikingly the assemblies grew much larger in the presence of RNA and NADH in seperate reactions (Figure 6B).

**Figure 6.**
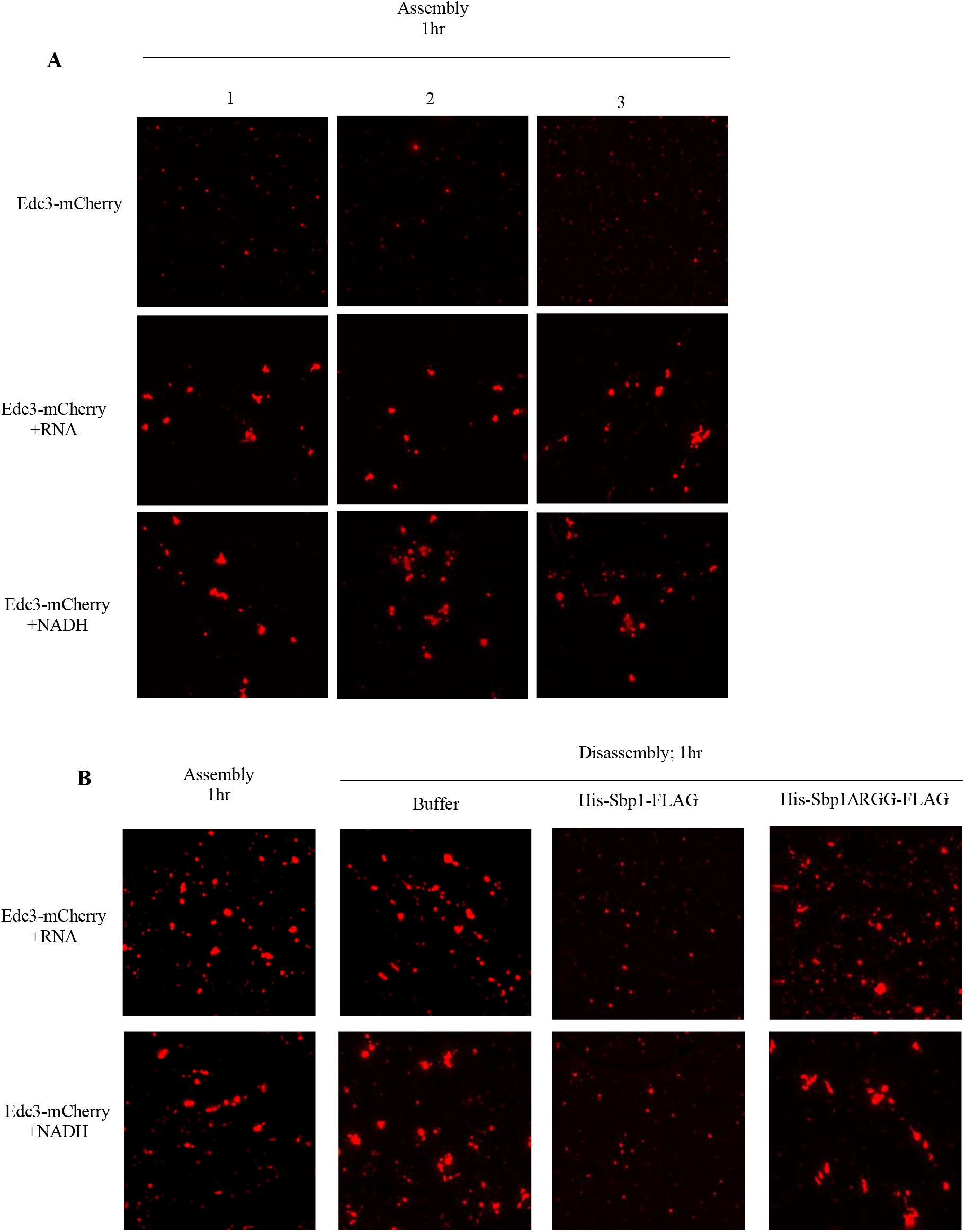
Purified Sbp1 disrupts Edc3 assemblies in RGG-motif dependent manner. A) Purified Edc3-mCherry (10µM) were kept at 30°C for 1 hour to phase-separate in LLPS buffer (150mM KCL; 30mM HEPES_KOH pH 7.4; 2mM MgCl_2_) in presence of RNA or NADH or both as indicated. 1, 2 and 3 indicate representative images from three independent experiments. B) After 1 hour the phase-separated assembly were than subjected to 30µM of Sbp1 or Sbp1ΔRGG protein and incubated again at 30°C for another 1 hour followed by microscopy at each time point during assembly and disassembly. The images were taken using DeltaVision DV Elite microscope.

Since the disassembly of Edc3 granules in cells was defective in the absence of Sbp1 and purified Sbp1 binds Edc3, we hypothesized that Sbp1 could dissolve Edc3 assemblies by physically interacting with it. Upon addition of purified Sbp1, we indeed observed reduction in the size of Edc3 assemblies. Decrease in the size of assemblies was dependent on the RGG-motif of Sbp1 as the addition of purified Sbp1Δrgg did not affect Edc3 assemblies (Figure 6B). We conclude based on these results that Edc3 can form higher order assemblies which is augmented by RNA and NADH. Further, these assemblies are dissovled upon incubation with purified Sbp1. This observation provides a mechanistic basis for the role of Sbp1 in P-body disassembly.

## Discussion

This study makes two important conclusions. It identifies a P-body disassembly factor in the form of Sbp1 and implicates a low complexity sequence (RGG-motif) in promoting disassembly. The following observations support these conclusions: 1) Edc3 granule disassembly is defective in the absence of Sbp1 upon recovery from oxidative and glucose starvation stress, 2) Disassembly of other P-body markers such as Dhh1 and Scd6 is also defective in the absence of Sbp1, 3) Complementation of *Δsbp1* strain by wild type *SBP1* but not sbp1Δrgg rescues the P-body disassembly defect, and 4) Purified Sbp1 but not Sbp1ΔRGG leads to decreased size of Edc3 assemblies *in vitro*.

Our data proposes a simple yet lucid model to explain the mechanistic basis of Sbp1 role in disassembly. Sbp1 directly binds Edc3 and this interaction abrogates the ability of Edc3 to form higher order assemblies. An obvious yet interesting question is the temporal regulation of this interaction. It may be hypothesized that the recruitment of Sbp1 to Edc3 assemblies may be promoted during recovery by a bridging adapter protein. It is also possible that Sbp1 and/or Edc3 undergo posttranslational modification(s) during recovery conditions that enable their interaction with each other. It is noteworthy that the deletion of Hmt1 (arginine methyltransferase) does not perturb the ability of Sbp1 to promote Edc3 disassembly (Supplementary Figure 3). This could be due to backup methyltransferase activity in the absence of Hmt1 as observed earlier (Bhatter et al., 2020). Importantly, the AMD (arginine methylation defective) mutant of Sbp1 fails to complement the Edc3 disassembly defect (Figure 4B & C) indicating that arginine methylation could be important for its role in disassembly.

It is intriguing that translation repressor protein Sbp1 itself localizes to P-bodies and also promotes its disassembly. It is currently unclear if the role of Sbp1 in repression and granule disassembly are related. It must be noted that both the functions of Sbp1 depend on its RGG-motif indicating that the two could be related.

Assembly of RNA granules during stress can be defective in the absence of translation repressors (Rajyaguru et al., 2012). Sbp1 localizes to RNA granules and its deletion decreases assembly of stress granule (Figure 1A & B). Since stress granules arise from pre-formed PB in yeast (Buchan et al., 2008), it is also likely that PB disassembly defect in *Δsbp1* could affect exchange of mRNPs contributing to the stress granule assembly defect. Understanding the connection between PB disassembly defect and stress granule assembly defect will be a future research direction.

Interestingly, we observe that NADH and RNA can independently augment formation of Edc3-mCherry assemblies (Figure 6). This is consistent with the report that *S. pombe* Edc3 phase separation is promoted by RNA (Damman et al., 2019). The IDR (FDF region) and the Yjef-N domain of Edc3 interact with RNA which facilitates phase separation. The role of NADH in Edc3 assembly formation has not been reported. It is known that human Edc3 binds NADH and both human and yeast Edc3 can chemically modify NAD (Walters et al., 2014). Mutants of yeast Edc3 predicted to be defective in binding NADH were defective in localizing to P-bodies (Walters et al., 2014). Our result provides a strong basis for focusing on the role of NADH in Edc3 function in vivo.

Previous reports suggest that autophagy plays an important role in stress granule clearance. VCP, a human ortholog of yeast Cdc48, is required for efficient stress granule clearance through granulophagy. Downregulation or chemical inhibition of VCP in HeLa cells led to a significant defect in stress granule clearance in mammalian cell culture (Buchan et al., 2013). ZFAND1 is a human ortholog of yeast Cuz1, a protein implicated in arsenite response in yeast (Hanna et al., 2014; Sa-Moura et al., 2013). ZFAND1 was reported to trigger proteasomal degradation of stress granule during arsenite-induced stress and its clearance during recovery. siRNA depletion of ZFAND1 resulted in granule clearance defect 2 h after recovery from arsenite stress (Turakhiya et al., 2018). Cuz1 deletion leads to defective SG clearance upon sodium arsenite treatment (cite Turakhiya) therefore we wondered if Cuz1 could affect P-body disassembly upon sodium azide stress. We observed that the assembly of stress granule and P-bodies were defective in *Δcuz1* as compared to wild-type (Supplementary Figure 4). However *Δcuz1* did not affect PB and SG disassembly (Supplementary Figure 4). This suggest that Cuz1 does not play a role in clearance of SG and PB that arise in response to sodium azide stress.

As mentioned in the introduction section, mutations in proteins with low complexity sequences such as FUS and TDP43 are involved in neurodegenerative disorders such ALS (Burke et al., 2015; Da Cruz and Cleveland, 2011; Murakami et al., 2015; Patel et al., 2015). These proteins form higher order assemblies which can be toxic (Sun et al., 2011). Evidence in literature suggest a role for Sbp1 in modulating FUS toxicity. Sbp1 was identified as as a multi-copy suppressor of FUS-related toxicity in yeast cells (Sun et al, 2011; Ju et al, 2011). The mechanistic basis of this observation is unclear. It is posible that Sbp1 could promote disassembly of FUS granules leading to suppression of toxicity which remains to be tested.

Overall our report identifies a bonafide PB disassembly factor which functions through low-complexity sequences. This report paves way to screen for and identify other disassembly factors of PB and RNA granules in general. Such factors will enhance our understanding of the role of RNA granules in regulation of mRNA fate under normal conditions and in diseases. Contrary to general understanding, our report brings to light a new role for LC sequences in granule disassembly.

## Materials and Methods

**Table 1:** List of strains used in this study.

|  |  |  |
| --- | --- | --- |
| yPIR1 | Wild-type | MATa *his3Δ1 leu2 met15 ura3* ('BY4741') |
| yPIR24 | $\Delta scd6$ | MATa *his3Δ1 leu2 ura3 his3 met15 scd6Δ::KanMX* |
| yPIR25 | $\Delta sbp1$ | MATa *his3Δ1 leu2 ura3 his3 met15 sbp1Δ::KanMX* |
| yPIR2 | $\Delta hmt1$ | MATa *his3Δ1 leu2Δ0 met15Δ0 ura3Δ0 hmt1Δ::KanMX* |
| yPIR82 | $\Delta cuz1$ | MATa *his3Δ1 leu2 ura3 his3 met15 cuz1Δ::KanMX* |

**Table 2:**
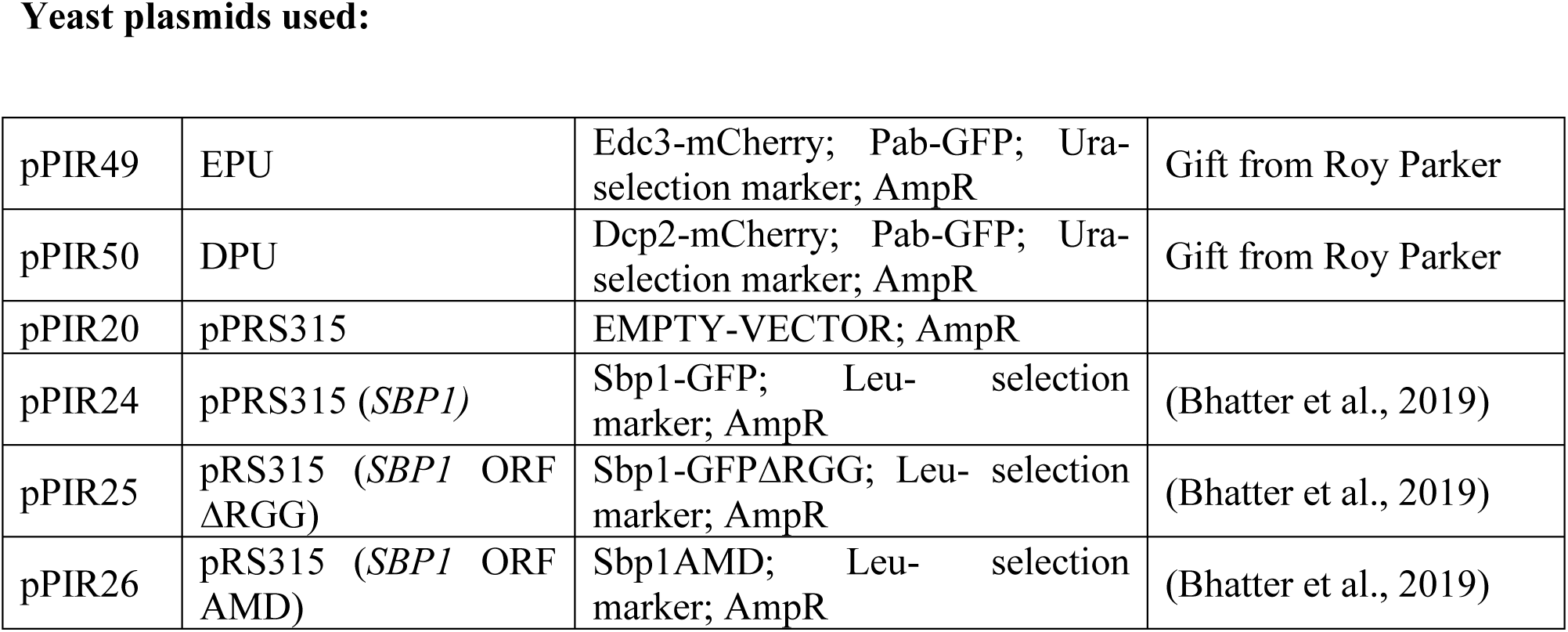
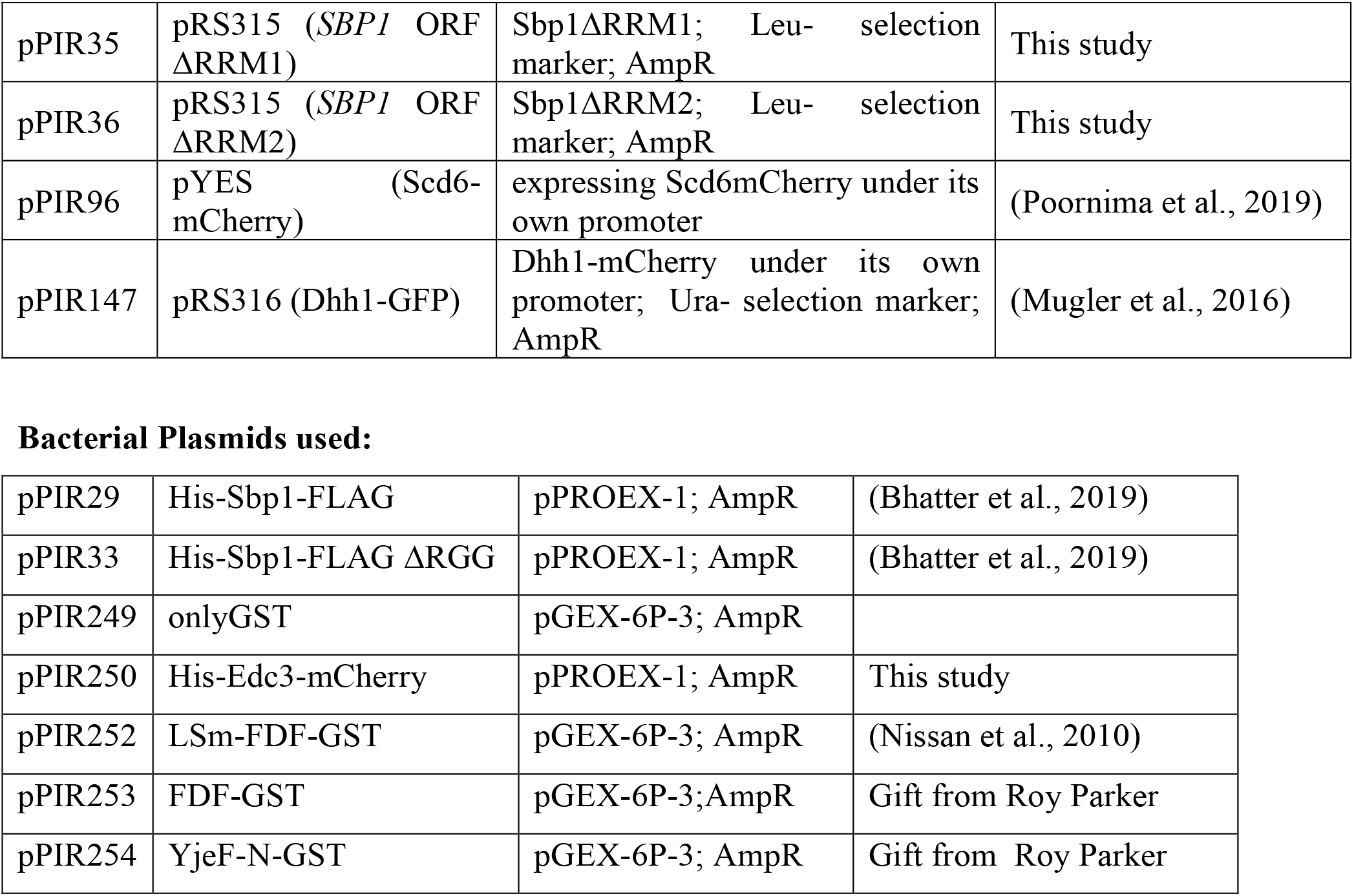
List of plasmids used in this study.

Yeast plasmids used:
|  |  |  |  |
| --- | --- | --- | --- |
| pPIR49 | EPU | Edc3-mCherry; Pab-GFP; Ura-selection marker; AmpR | Gift from Roy Parker |
| pPIR50 | DPU | Dcp2-mCherry; Pab-GFP; Ura-selection marker; AmpR | Gift from Roy Parker |
| pPIR20 | pPRS315 | EMPTY-VECTOR; AmpR |  |
| pPIR24 | pPRS315 (*SBP1*) | Sbp1-GFP; Leu-selection marker; AmpR | (Bhatter et al., 2019) |
| pPIR25 | pRS315 (*SBP1* ORF ΔRGG) | Sbp1-GFPΔRGG; Leu-selection marker; AmpR | (Bhatter et al., 2019) |
| pPIR26 | pRS315 (*SBP1* ORF AMD) | Sbp1AMD; Leu-selection marker; AmpR | (Bhatter et al., 2019) |
| pPIR35 | pRS315 (*SBP1* ORF ΔRRM1) | Sbp1ΔRRM1; Leu- selection marker; AmpR | This study |
| pPIR36 | pRS315 (*SBP1* ORF ΔRRM2) | Sbp1ΔRRM2; Leu- selection marker; AmpR | This study |
| pPIR96 | pYES (Scd6-mCherry) | expressing Scd6mCherry under its own promoter | (Poornima et al., 2019) |
| pPIR147 | pRS316 (Dhh1-GFP) | Dhh1-mCherry under its own promoter; Ura- selection marker; AmpR | (Mugler et al., 2016) |
| Bacterial Plasmids used: |  |  |  |
| pPIR29 | His-Sbp1-FLAG | pPROEX-1; AmpR | (Bhatter et al., 2019) |
| pPIR33 | His-Sbp1-FLAG ΔRGG | pPROEX-1; AmpR | (Bhatter et al., 2019) |
| pPIR249 | onlyGST | pGEX-6P-3; AmpR |  |
| pPIR250 | His-Edc3-mCherry | pPROEX-1; AmpR | This study |
| pPIR252 | LSm-FDF-GST | pGEX-6P-3; AmpR | (Nissan et al., 2010) |
| pPIR253 | FDF-GST | pGEX-6P-3;AmpR | Gift from Roy Parker |
| pPIR254 | YjeF-N-GST | pGEX-6P-3; AmpR | Gift from Roy Parker |

**Table 3:**
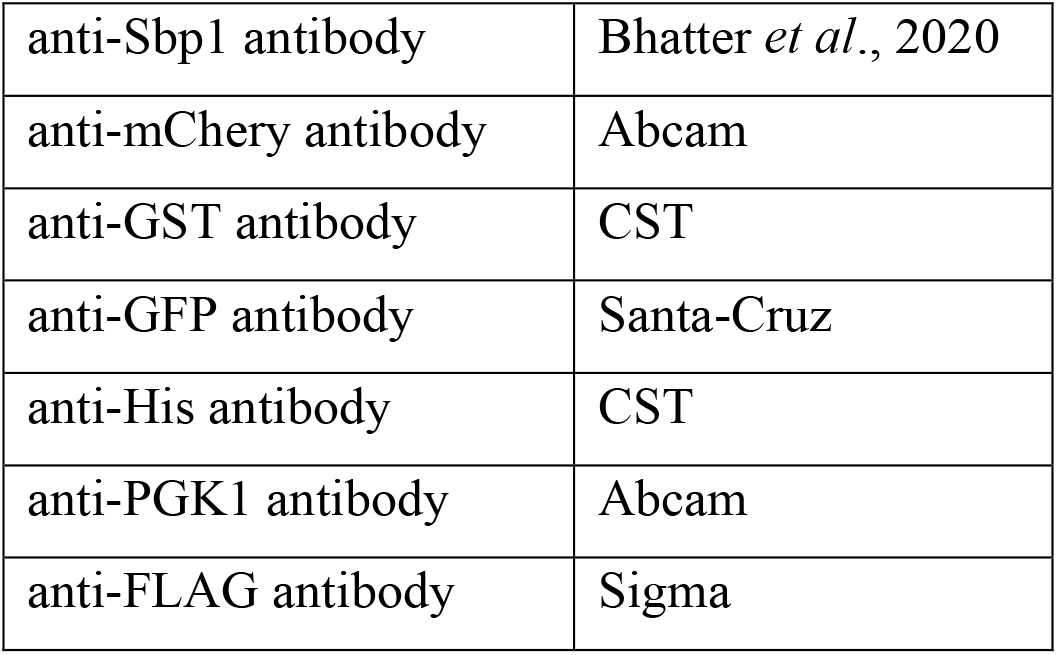
List of antibodies used in this study.

### Media and cultures

Overnight culture of yeast strains (transformed with the indicated plasmid) grown in SD Ura-yeast growth medium was diluted to OD_600_=0.1 and grown at 30°C until they reached an OD_600_ of 0.4-0.5. Cultures were then incubated for 30 min with 0.5% sodium azide v/v at 30°C or with equal volume water. Subsequently, the cells were pelleted by centrifugation (4200g, 10 seconds, RT) and washed twice with glucose-containing SD ura-medium. For stress recovery, the resuspended cells were grown for an additional 1 hour at 30°C in media without sodium azide. 1.5 mL cells were collected by centrifugation (14000g, 12 sec, RT), supernatant was removed and the cells were resuspended in 20 ul of fresh media or residual media and were spotted on coverslips for microscopy.

For glucose starvation experiment, overnight cultures of yeast strains (transformed with the indicated plasmid) in SD Ura-yeast growth medium was diluted to OD_600_=0.1 and grown at 30°C until they reached an OD_600_ of 0.4 to 0.5. Cultures were divided into two equal part and washed one part with glucose-containing media and the other part with glucose deprived media. Cultures were then incubated for 30 min in the presence and absence of 2% glucose at 30°C. Subsequently, the cells were pelleted by centrifugation (4200g, 10 seconds, RT) and washed twice with SD ura-medium with/without glucose. For stress recovery, the resuspended cells were grown for an additional 1 hour at 30°C. Microscopy was carried out in each of the conditions mentioned above as depicted in figure 3. For microscopy, 1.5 mL cells were collected by centrifugation (14000g, 12 sec, RT). Supernatant was removed, and the cells were resuspended, then immobilized on coverslips for microscopy.

### Microscopy

All images were acquired using a Deltavision RT microscope system running softWoRx 3.5.1 software (Applied Precision, LLC), using an Olympus 100×, oil-immersion 1.516 NA objective. Exposure time and transmittance settings for Green Fluorescent Protein (GFP) channel were 0.2 seconds and 32%, and for mCherry channel were 0.3 seconds and 32% respectively. Images were collected as 512×512-pixel files with a CoolSnapHQ camera (Photometrics) using 2×2 binning for yeast. All yeast images were deconvolved using standard softWoRx deconvolution algorithms for granule counting and data analysis. For each experiment, more than 100 cells were considered for granule counting manually Data from three independent experiments was used for quantitation and statistical significance was calculated using non-parametric student t-test.

### Complementation experiment

For complementation experiment we constructed a CEN plasmid (pRS315) expressing wild-type Sbp1 and its mutants (Sbp1ΔRGG, Sbp1ΔRRM1, Sbp1ΔRRM2 and Sbp1-AMD). *SBP1* ORF was amplified using specific primers from *S. cerevisiae* genomic DNA along with its promoter. The plasmid was digested with SmaI resulting in blunt end plasmid DNA. The amplified SBP1 was then ligated to the digested pRS315 plasmid using T4 DNA ligase (Thermo). The mutants were created by the same method for site-directed mutagenesis using specific primers designed from Agilent QuickChange Primer design web tool. The *SBP1* gene containing plasmid construct and its mutants was transformed in *Δsbp1 and* wild-type strain along with empty-vector as control followed by live-cell imaging.

### Western analysis

For performing western to estimate Edc3-mCherry levels in wild-type and *Δsbp1* under unstressed, stressed and recovery conditions, overnight cultures were diluted to 0.1 O.D. and grown till 0.4-0.5 O.D. Cultures were then incubated for 30 min with 0.5% sodium azide v/v at 30°C or without sodium azide. Subsequently, the cells were pelleted by centrifugation (4200g, 1 minute, RT) and washed twice with glucose-containing SD ura-medium. For stress recovery, the resuspended cells were grown for an additional 1 hour at 30°C. Unstressed, stressed and recovery cells were collected in a 50 mL falcon by centrifuging at 4200g for 1 min. Cells were then broken open in 100 µl lysis buffer containing 50 mM Tris–Cl pH7.5, 50 mM NaCl, 2 mM MgCl2, 0.1% Triton-X100, 1 mM β-Mercaptoethanol, 1× Complete mini-EDTA-free tablet (Roche, catalog no. 04693132001) and lysed by vortexing at 4°C in bead-beater with glass beads. Unbroken cells and debris were removed by centrifugation at 5500 rpm for 5 min at 4°C, followed by a 1 min spin at 13000 rpm to remove any protein aggregates. A total of 100 µg of total protein was used for loading onto SDS-PAGE. Western analysis was performed using anti-GST (CST, catalog no. 2624; 1:1000 dilution), anti-His (CST, catalog no. 2366; 1:1000 dilution), anti-GFP (Santa Cruz, catalog no. sc-9996; 1:1000 dilution), and anti-PGK1 (Abcam, catalog no. ab113687; dilution 1:1000.

### Protein expression and purification

For *in vitro* binding assay, proteins were purified from E. coli BL21 by batch purification using glutathione sepharose (GE Healthcare, Chicago, IL, USA, catalog no. 17075604) or Ni-NTA agarose (ThermoFisher Scientific, catalog no. 88222). Cell pellet was resuspended in lysis buffer along with addition of lysozyme (10 μg·mL−1), DTT (1 mm), PMSF (2 mm) and Protease Inhibitor Tablet (Roche, Basel, Switzerland), RNase A (1 mg·mL−1) for 20 min. Lysis was done using sonication. And debris were separated by centrifuging at 15000 rpm for 15 minutes at 4°C. Lysate was allowed to bind to beads for 1 h on nutator at 4 °C. Washes were performed on nutator for 10 min at 4° with Ni-NTA wash buffer (300mM NaCl, 150 mM NaH_2_PO_4_, 20-40mM imidazole) for His-tagged purification and with 1x PBS for GST-tagged purification. For His tagged proteins, elution was done with 250 mm of imidazole (SRL cat# 61510) while for GST tagged proteins were kept bead bound in storage buffer (10mM Tris Base pH 7.0, 25mM NaCl, 1mM DTT, 20% Glycerol) at 4°C for immediate use. Purified his-tagged protein was concentrated and dialyzed into 10 mm Tris–Cl pH7.5, 100 mm NaCl, 20% glycerol and 1 mm DTT in the cold room. Western analysis was performed using anti-GST (CST, catalog no. 2624; 1:1000 dilution), anti-Sbp1 antibody (home-made) and anti-FLAG antibody(Sigma) for checking purity of protein and then used for binding studies.

For Edc3 assembly assay, proteins were purified from *E. coli* BL21 by batch purification using Ni-NTA agarose (ThermoFisher Scientific, catalog no. 88222). Cell pellet was resuspended in L-arginine lysis buffer (50mM Tris Cl H 7.4; 500mM NaCl; 10% Glycerol; 5mM β-mercaptoethanol; 200mM L-arginine pH 7.4) along with the addition of lysozyme (10 μg·mL−1), DTT (1 mm), PMSF (2 mm) and Protease Inhibitor Tablet (Roche, Basel, Switzerland), RNase A (1 mg·mL−1) for 20 min. Lysis was done using sonication (SONICS, VibraCell; Pulse rate of 10 sec ON and 10 sec OFF for 1:30 minutes). Debris was separated by centrifuging at 15000 rpm for 15 minutes at 4°C. Lysate was allowed to bind to beads overnight on nutator at 4 °C. Washes were performed on nutator for 10 min at 4° with Ni-NTA wash buffer (300mM NaCl, 150 mM NaH_2_PO_4_, 20-40mM imidazole). Elution was done with 250 mm of imidazole (SRL cat# 61510). Purified his-tagged protein was concentrated and dialyzed into 10 mm Tris–Cl pH7.5, 100 mm NaCl, 20% glycerol and 1 mm DTT in the cold room. Western blot analysis was done with anti-mCherry antibody (Abcam) to check the purity of the purified protein before carrying out liquid-droplet experiment.

### *In vitro* pulldown

For recombinant protein pull-downs, 200 pmoles of purified proteins were incubated with immobilized GST-tagged protein to glutathione sepharose beads (GE Healthcare) at 4°C for binding reactions (for 2h). The binding buffer for Glutathione pull downs contained 50Mm HEPES pH7, 100mM NaCl, 1mM DTT, 2mM MnCl_2_, 2mM MgCl_2_, 1% Triton-X100, 10% glycerol, 0.25mg/ml RNase A and 10mg/ml BSA. The beads were washed thrice with binding buffer, 10μl of SDS-PAGE loading dye was added to beads and analyzed by SDS-PAGE followed by Western blotting.

### Edc3 assembly assay

Reactions were performed in 1.5 mL micro-centrifug tube. Proteins were diluted to 10µM in liquid droplet buffer containing 150 mM KCl, 30 mM HEPES-KOH pH 7.4, 2 mM MgCl2. 1µg of total RNA or 0.2mM NADH or both were added in the reaction of 30µL and kept at 30°C untouched for liquid-droplet formation for 1 hour. The reactions were centrifuged at 200g for 2 minutes and supernatant was discarded very gently leaving 10uL of droplet reaction followed by microscopy.

For the droplet disassembly experiment, Sbp1, Sbp1ΔRGG, BSA or buffer were added respectively in the above preformed droplet reactions after 1 hour and incubated for 1 hour more at 30°C. The reactions were centrifuged at 200g for 2 minutes and supernatant was discarded very gently leaving 10uL of droplet reaction followed by microscopy. Microscopy was performed using a Deltavision RT microscope system running softWoRx 3.5.1 software (Applied Precision, LLC), using an Olympus 100×, oil-immersion 1.516 NA objective. Exposure time and transmittance settings for Green Fluorescent Protein (GFP) channel were 0.2 seconds and 32%, and for mCherry channel were 0.3 seconds and 32% respectively. Images were collected as 512×512-pixel files with a CoolSnapHQ camera (Photometrics) using 2×2 binning for yeast. All yeast images were deconvolved using standard softWoRx deconvolution algorithms for granule counting and data analysis.

## Acknowledgements

We are indebted to Roy Parker for providing us various plasmids. We also thank Karsten Weiss for his generous gift of Dhh1-GFP plasmid.

## Author contributions

Conceptualization and hypothesis-PIR and RR; Experimental design-PIR and RR; Experimentation-RR and IAK; Data interpretation-PIR, RR and IAK; Manuscript writing (first draft) – PIR, Subsequent draft review and editing – PIR and RR; NB created the pRS315 (*SBP1ΔRRM1*) and pRS315(*SBP1ΔRRM2*) mutant constructs from pRS315(*SBP1)* created by RR.

## Notes

### Competing Interest Statement

The authors have declared no competing interest.

