## Supplementary figures for "RGG-motif protein Sbp1 is required for Processing body (P-body) disassembly"

**Supplementary Figure 1**

**B**

**A**

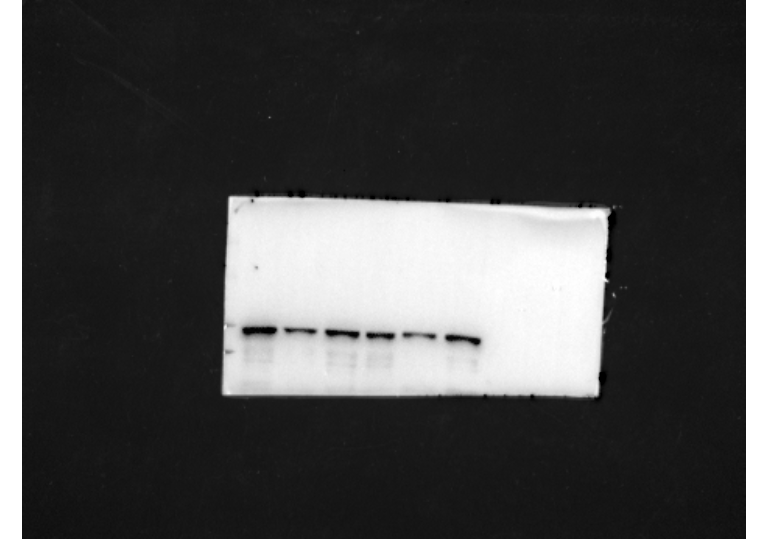

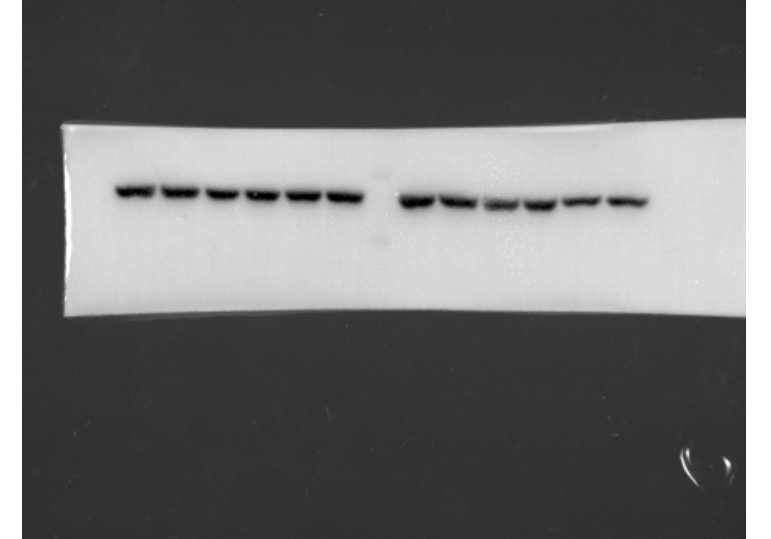

Wild-type –Sodium azide

Wild-type + Sodium azide

*Δsbp1+*Sodium azide

*Δsbp1* recovery

*Δsbp1–*Sodium azide

Wild-type recovery

anti-mCherry

anti-PGK1

Edc3-mCherry

PGK1

100

75

50

37

kDa

Level of Edc3-mCherry protein

Wild-type

*Δsbp1*

Recovery

-Sodium azide

+Sodium azide

Recovery

-Sodium azide

+Sodium azide

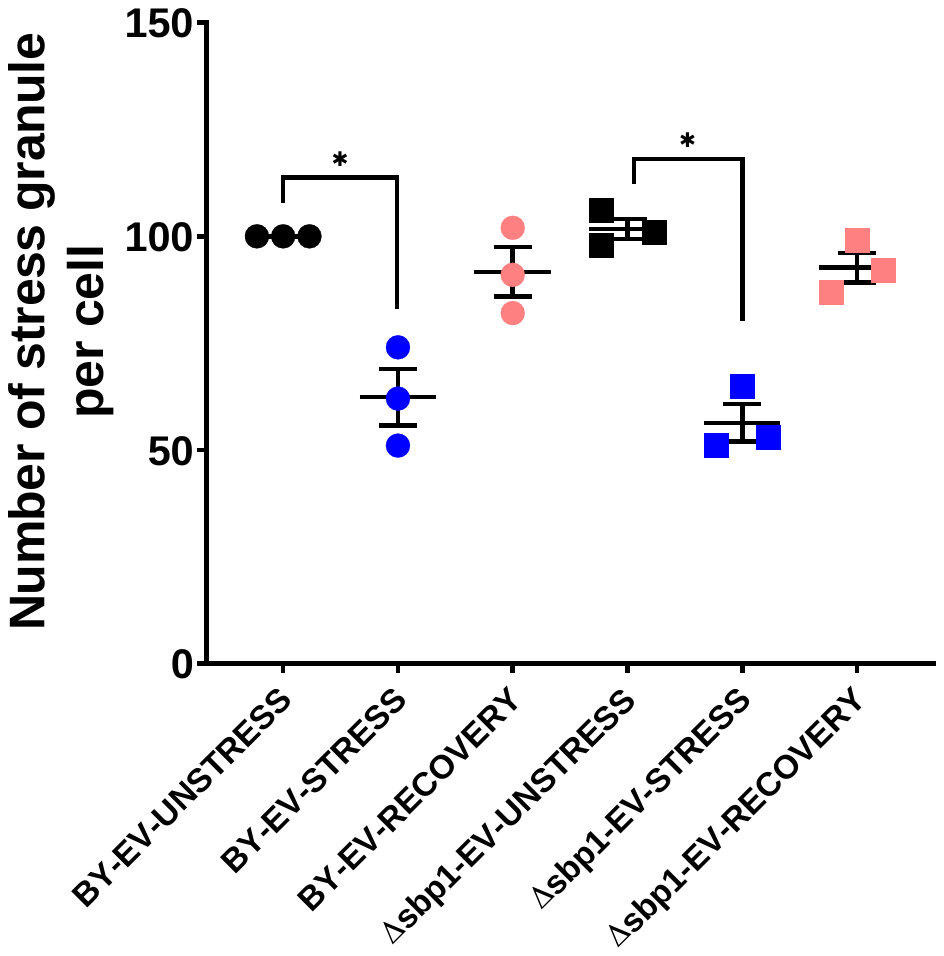

**Supplementary Figure 1. Edc3-mCherry granule disassembly defect is not stress specific**

A) Protein levels of Edc3-mCherry does not change in *Δsbp1* as compared to wild-type during recovery from glucose starvation stress. B) Quantification of Edc3-mCherry protein level from the glucose starvation stress experiment done in A (p<0.005).

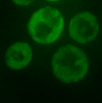

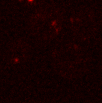

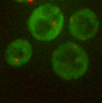

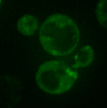

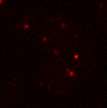

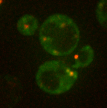

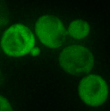

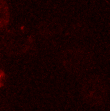

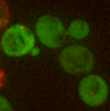

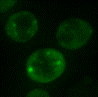

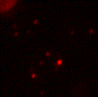

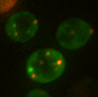

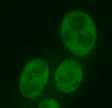

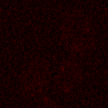

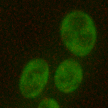

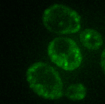

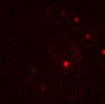

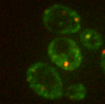

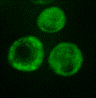

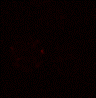

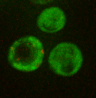

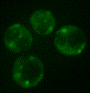

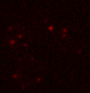

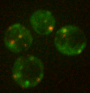

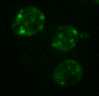

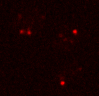

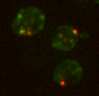

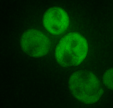

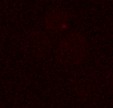

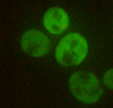
**Supplementary Figure 2**

*Δsbp1-*pRS315 (EV)

-Sodium azide

+Sodium azide

*Δsbp1-*pRS315 (*SBP1*)

*Δsbp1-*pRS315 *(SBP1ΔRGG)*

-Sodium azide

+Sodium azide

*Δsbp1-*pRS315 *(SBP1-AMD)*

Pab1-GFP

Edc3-mCherry

Merge

-Sodium azide

+Sodium azide

Pab1-GFP

Edc3-mCherry

Merge

Wild-type pRS315 (EV)

**A**

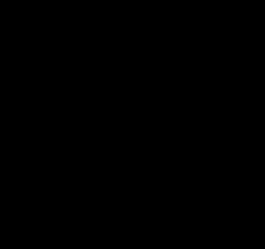

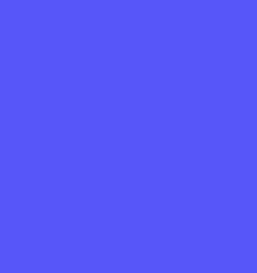

+Sodium azide

-Sodium azide

**B**

BY4741-pRS315 (EV)

*Δsbp1-*pRS315 *(EV)*

*Δsbp1-*pRS315 *(SBP1)*

*Δsbp1-*pRS315 *(SBP1ΔRGG)*

*Δsbp1*-pRS315 *(SBP1-AMD)*

Number of Pab1-GFP granule per cell

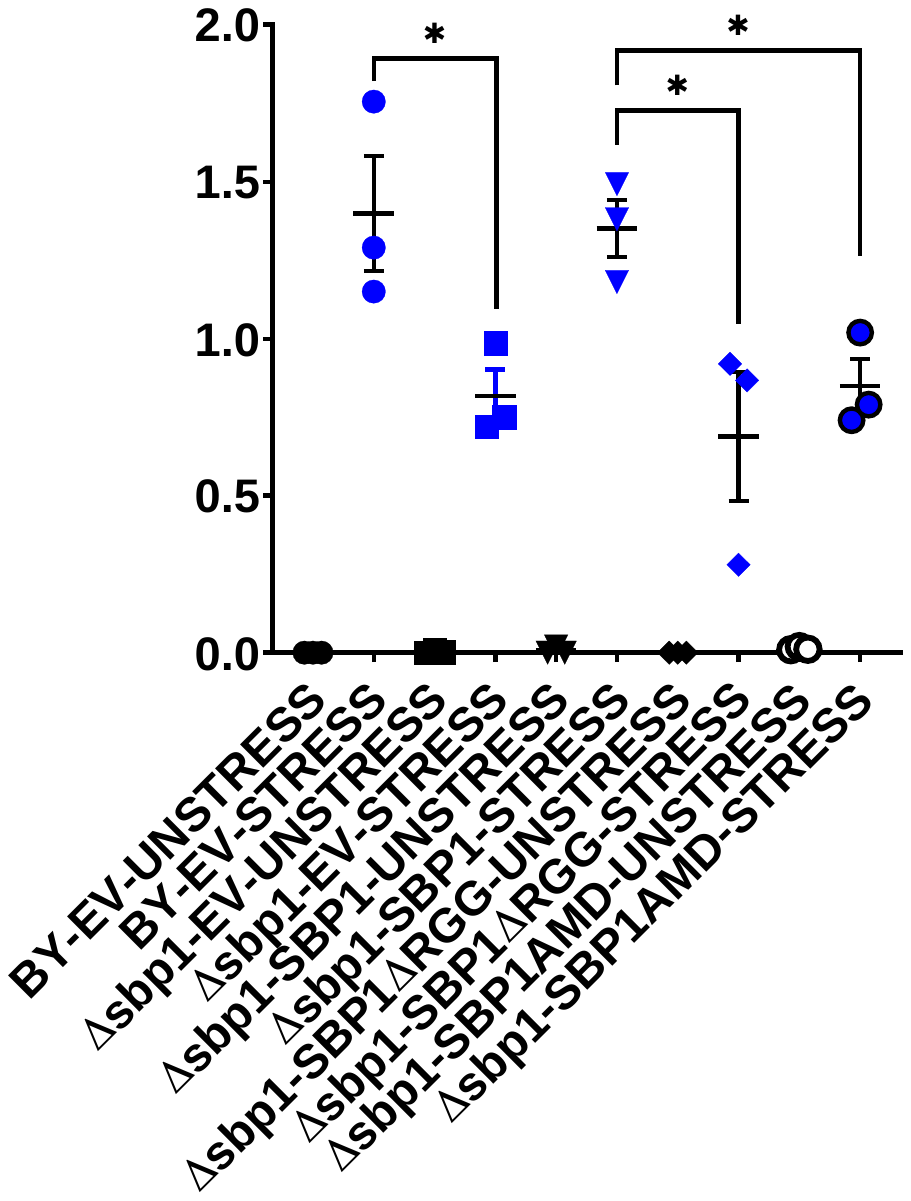

BY4741-pRS315 (EV)

*Δsbp1-*pRS315 *(EV)*

*Δsbp1*-pRS315 *(SBP1)*

*Δsbp1-*pRS315 *(SBP1ΔRGG)*

*Δsbp1-*pRS315 *(SBP1-AMD)*

Number of Edc3-mCherry granule per cell

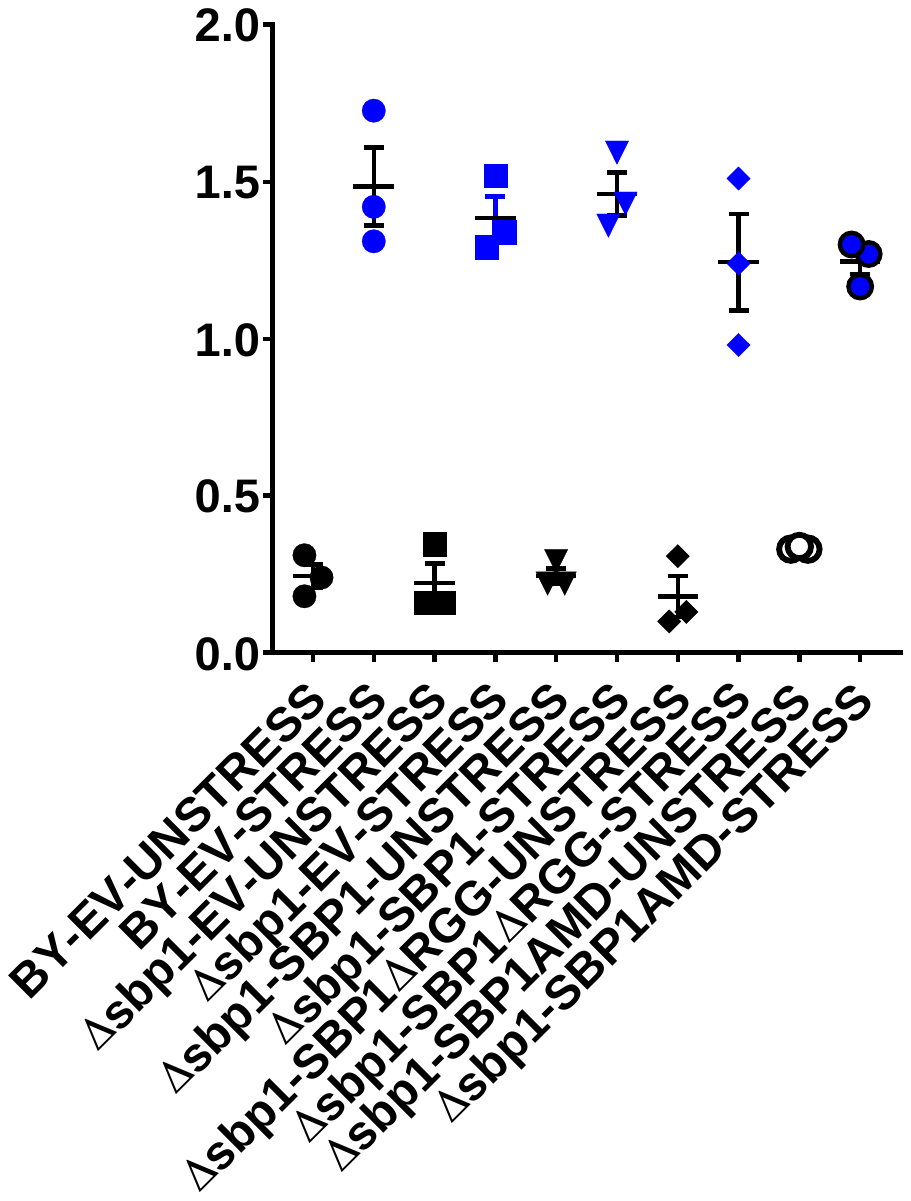

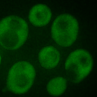

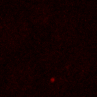

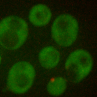

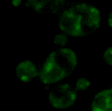

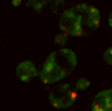

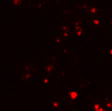

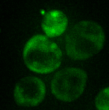

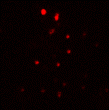

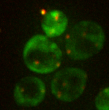

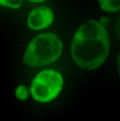

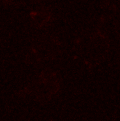

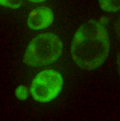

*Δsbp1-*pRS315 *(SBP1ΔRRM1)*

*Δsbp1-*pRS315 *(SBP1ΔRRM2)*

Pab1-GFP

Edc3-mCherry

Merge

Pab1-GFP

Edc3-mCherry

Merge

**C**

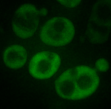

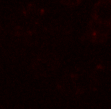

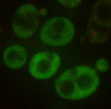

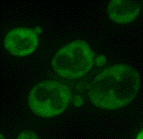

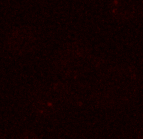

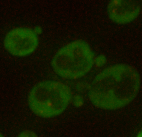

**D**

Number of Edc3-mCherry granule per cell

Number of Pab1-GFP granule per cell

*Δsbp1-*pRS315 *(SBP1)*

*Δsbp1-*pRS315 *(SBP1ΔRRM1)*

*Δsbp1-*pRS315 *(SBP1ΔRRM2)*

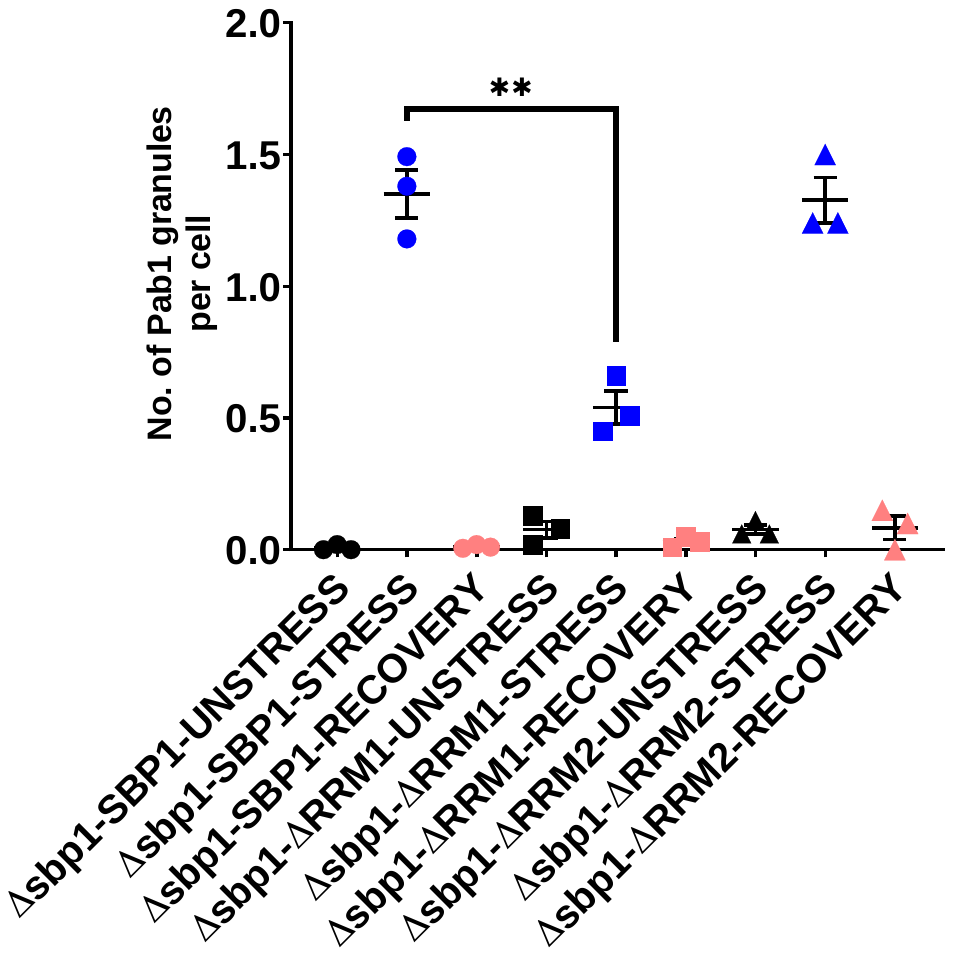

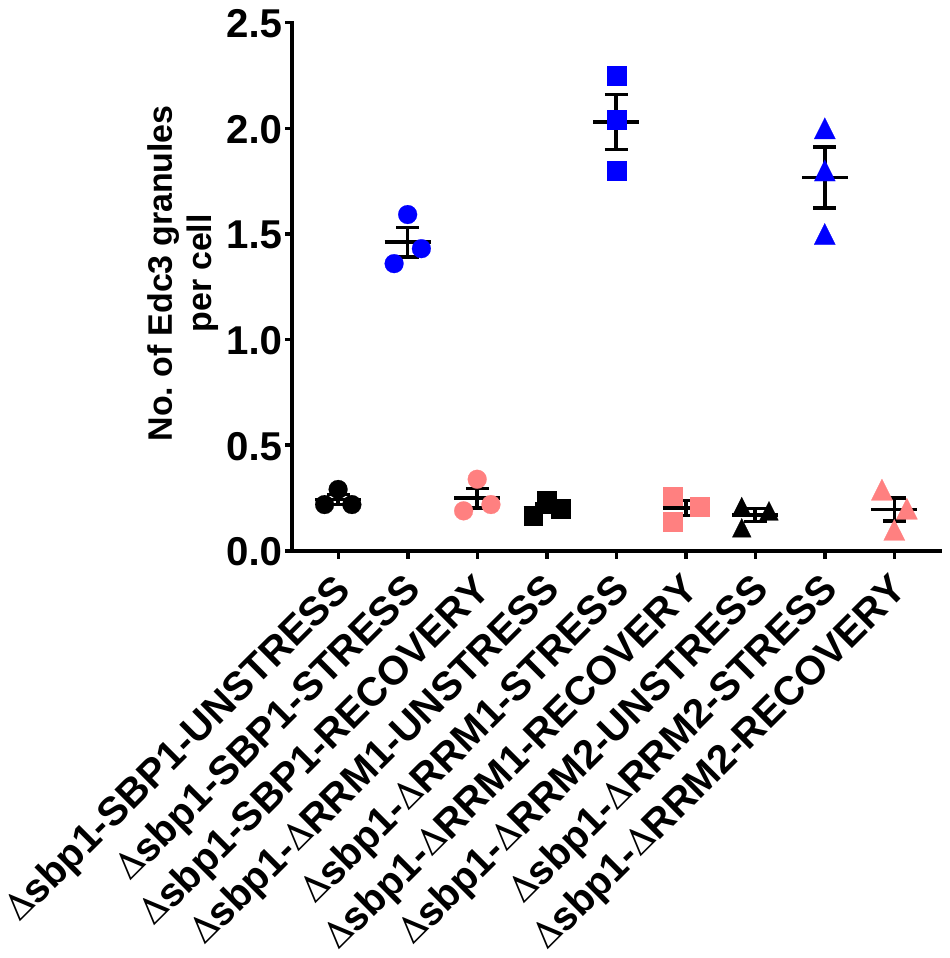

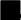

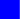

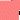

–sodium azide

+sodium azide

Recovery

*Δsbp1-*pRS315 *(SBP1)*

*Δsbp1-*pRS315 *(SBP1ΔRRM1)*

*Δsbp1*-pRS315 *(SBP1ΔRRM2)*

**Supplementary Figure 2: Complementation of *Δsbp1* with wild-type *SBP1* and mutants.**

A) Live-cell microscopy during unstress and stress condition for the recovery experiment. Plasmid containing wild-type Sbp1 and Sbp1 mutants were transformed in *Δsbp1* cells as indicated. Cells were cultured till 0.35-0.4 O.D. and incubated for 30 minutes with or without 0.5% (v/v) Sodium azide at 30℃. B) Quantitation of the complementation experiment described in A (p<0.005). C) Live-cell microscopy during unstress and stress condition for the recovery experiment. Plasmid containing wild-type Sbp1 and Sbp1 mutants (*ΔRRM1 and ΔRRM2)* were transformed in *Δsbp1*cells as indicated. Cells were cultured till 0.35-0.4 O.D. and incubated for 30 minutes with or without 0.5% (v/v) Sodium azide at 30℃. D) Quantitation of the complementation experiment described in C (p<0.005).

**Supplementary Figure 3**

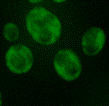

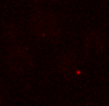

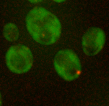

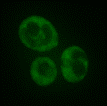

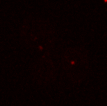

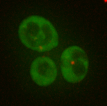

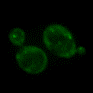

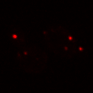

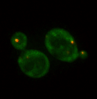

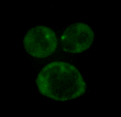

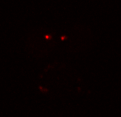

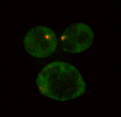

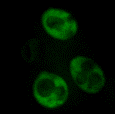

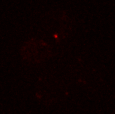

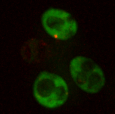

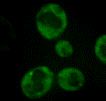

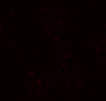

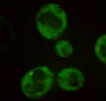

Recovery

*Δsbp1*

Wild-type

*Δhmt1*

Wild-type

*Δhmt1*

Wild-type

*Δhmt1*

+Sodium azide

-Sodium azide

**A**

Wild-type

*Δhmt1*

Wild-type

*Δhmt1*

Wild-type

*Δhmt1*

Recovery

-Sodium azide

+Sodium azide

Number of Pab1-GFP granule per cell

**B**

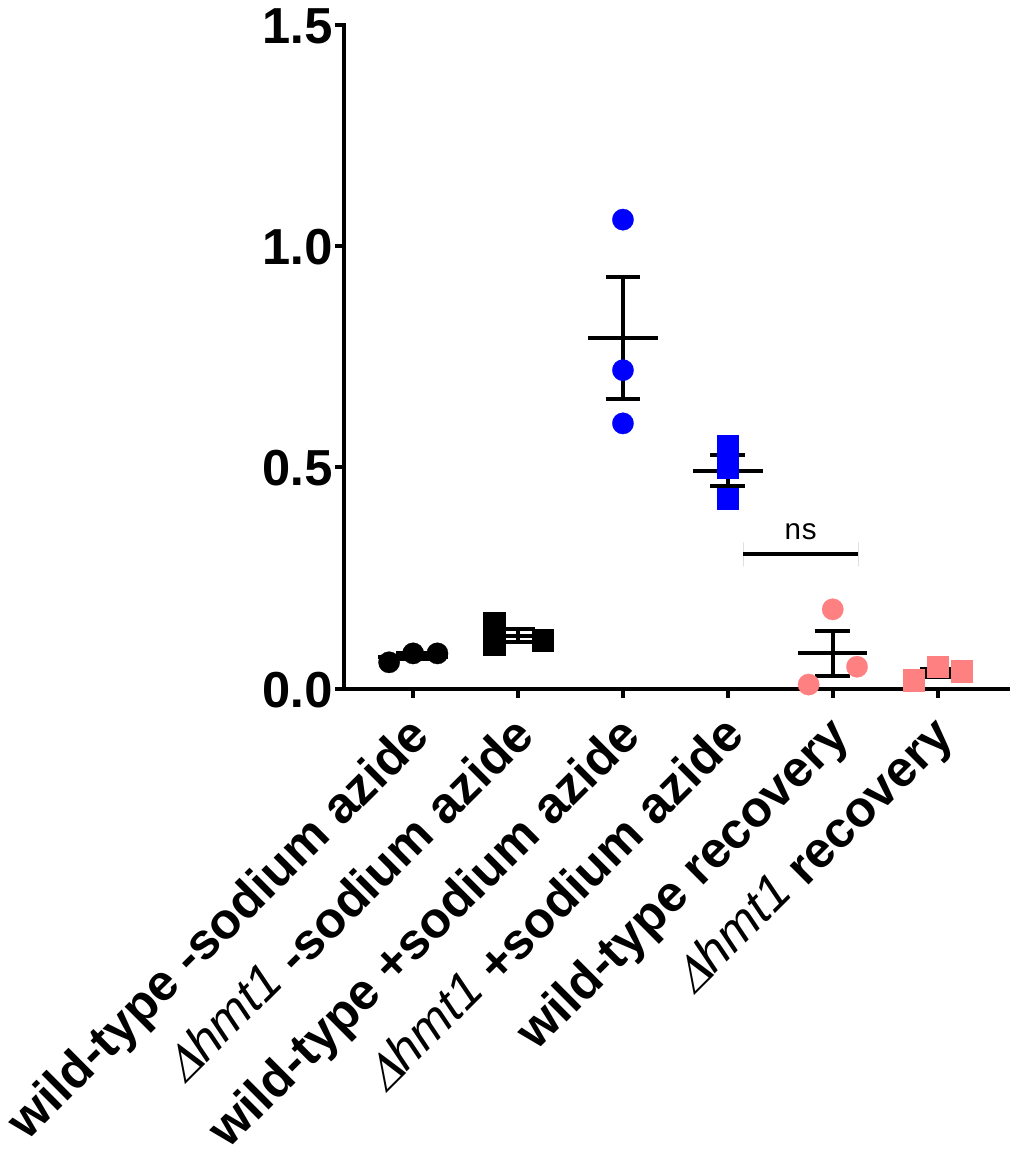

Wild-type

*Δhmt1*

Wild-type

*Δhmt1*

Wild-type

*Δhmt1*

Recovery

-Sodium azide

+Sodium azide

Number of Edc3-mCherry granule per cell

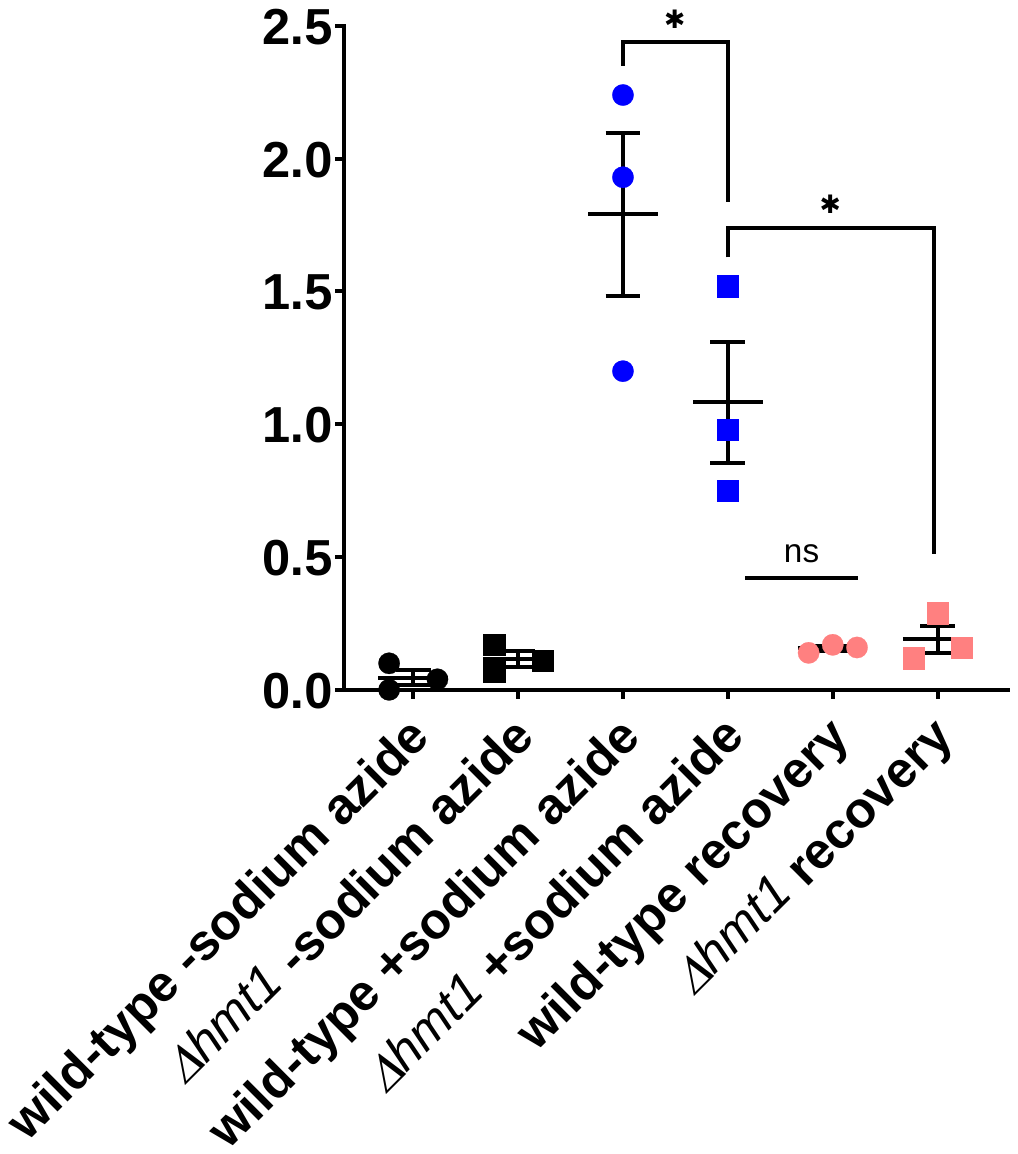

**Supplementary Figure 3. Hmt1 deletion does not lead to Edc3 granule disassembly defect.**

A) *Δhmt1 c*ells were cultured till 0.35-0.4 O.D. and incubated for 30 minutes with or without 0.5% (v/v) sodium azide at 30℃. Subsequently, cells were pelleted by centrifugation (4200g, 10 seconds, RT) and washed thrice with glucose containing medium. For stress recovery, the resuspended cells were grown for an additional 1 hour at 30℃ in media without sodium azide. B) Graph depicting the number foci per cell in wild-type and *Δhmt1* strain in various culture conditions (p<0.005).

**Supplementary Figure 4**

-Sodium azide

+Sodium azide

Recovery

Number of Edc3-mCherry granule per cell

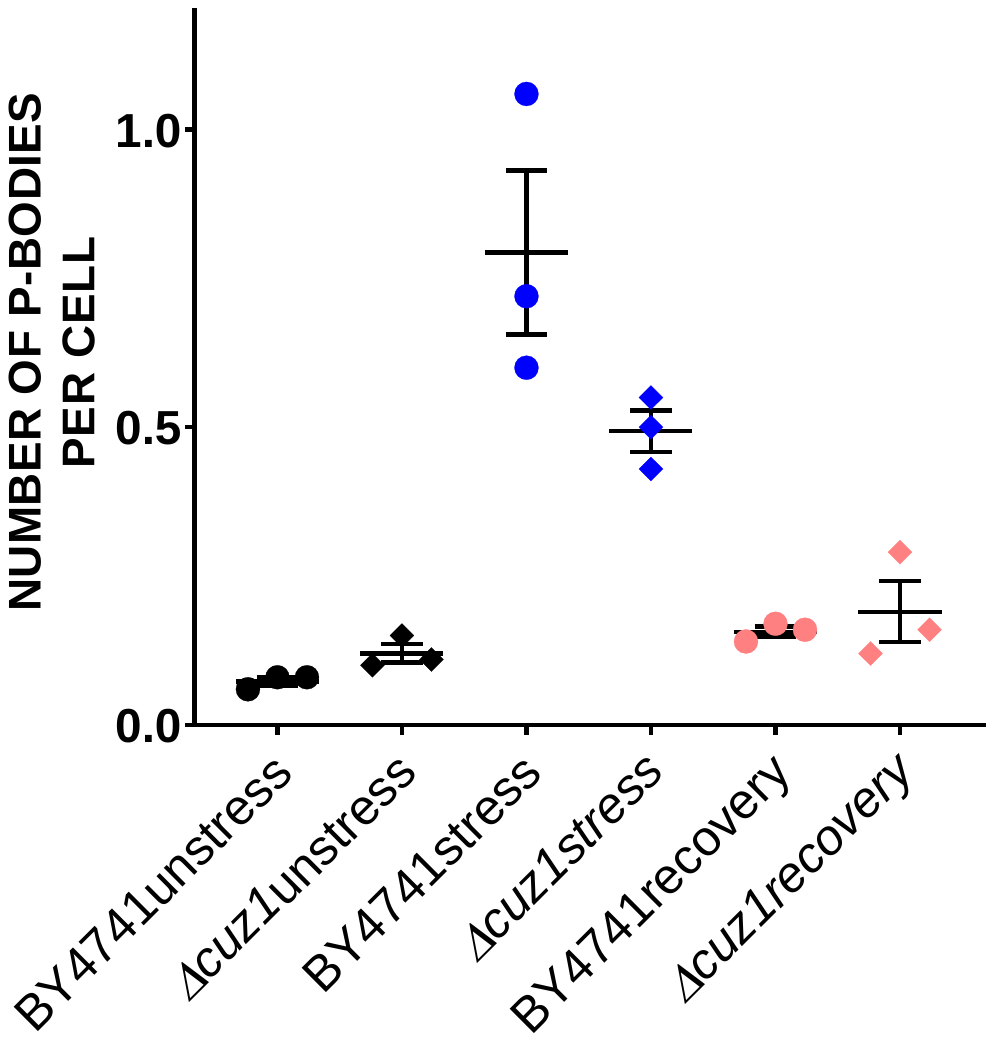

Wild-type

*Δcuz1*

Wild-type

*Δcuz1*

Wild-type

*Δcuz1*

**B**

Number of Pab1-GFP granule per cell

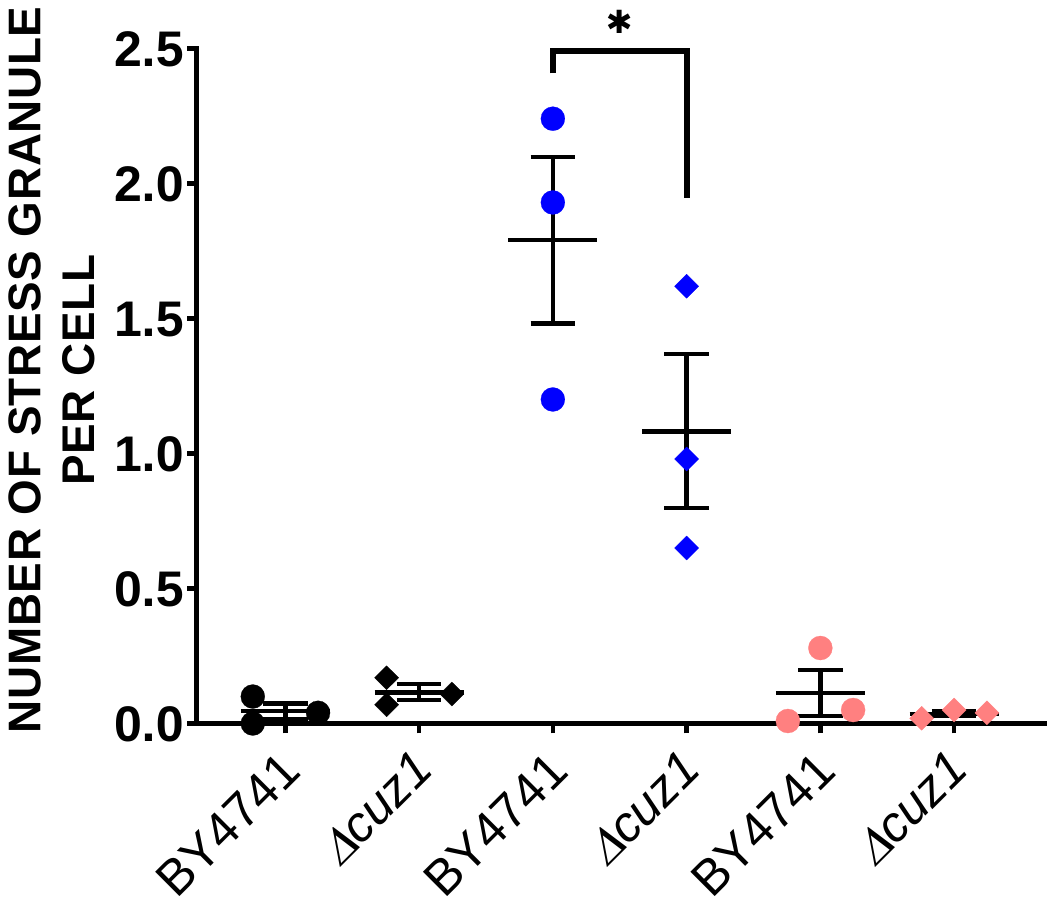

-Sodium azide

+Sodium azide

Recovery

Wild-type

*Δcuz1*

Wild-type

*Δcuz1*

Wild-type

*Δcuz1*

**+Sodium azide**

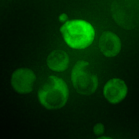

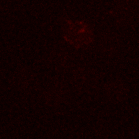

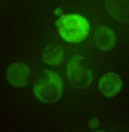

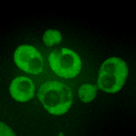

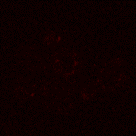

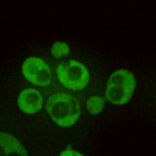

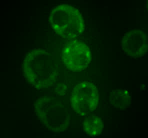

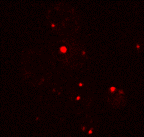

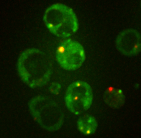

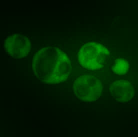

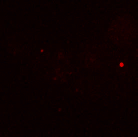

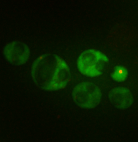

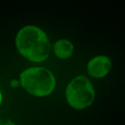

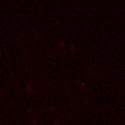

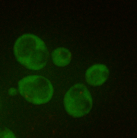

**Recovery**

**Pab1-GFP**

**Edc3-mCherry**

**Merged**

**Wild-type**

***Δcuz1***

**A**

**Wild-type**

***Δcuz1***

**Wild-type**

***Δcuz1***

**-Sodium**

**azide**

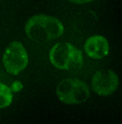

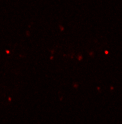

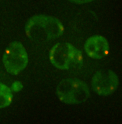

**Supplementary Figure 4: Cuz1 deletion does not lead to Edc3 granule disassembly defect.**

A) *Δcuz1 c*ells were cultured till 0.35-0.4 O.D. and incubated for 30 minutes with or without 0.5% (v/v) sodium azide at 30℃. Subsequently, cells were pelleted by centrifugation (4200g, 10 seconds, RT) and washed thrice with glucose containing medium. For stress recovery, the resuspended cells were grown foe an additional 1 hour at 30℃ in media without sodium azide. B) Graph depicting the number foci per cell in wild-type and *Δcuz1* strain in various culture conditions (p<0.005).
